# Hydrogen sulfide protects erectile function through stimulation of antioxidant defense in the corpus cavernosum

**DOI:** 10.1101/2025.05.05.652305

**Authors:** Tooyib A. Azeez, Paige R. Rhein, Clifford J. Pierre, Colin M. Ihrig, Justin D. La Favor

## Abstract

Obesity is a major risk factor for erectile dysfunction, whereby excess reactive oxygen species (ROS) in the corpus cavernosum have been implicated as a causative factor. Hydrogen sulfide (H_2_S) is an endogenous gasotransmitter with vasodilatory and antioxidant properties. The objectives of this study were to determine the influence of H_2_S on erectile function, penile ROS, and expressions of endogenous antioxidant genes and proteins in the corpus cavernosum. These objectives were tested by 1) comparing wild type mice to mice deficient in cystathionine γ-lyase (CSE^-/-^), a major endogenous source of H_2_S; and 2) by feeding wild type mice a control diet or high-fat, high-sucrose Western style diet (WD) for 18 weeks, with subgroups treated with and without the H_2_S prodrug SG1002 for the final 6 weeks of the dietary intervention. Erectile function was assessed by cavernous nerve-stimulated intracavernous pressure measurement, penile interstitial ROS was measured with a microdialysis approach, and cavernous gene and protein expression assessed by qRT-PCR and western blot, respectively. Erectile function was impaired in CSE^-/-^ mice and following the WD, which was significantly improved in WD mice treated with SG1002. Penile ROS levels were increased in CSE^-/-^ mice and following the WD, which were suppressed in WD mice treated with SG1002. The following genes were commonly decreased in CSE^-/-^ mice and increased by SG1002 treatment: Nqo1, Gclc, Gstm1, Gpx1, Gpx4, Hmox1, Txnrd1, and Prdx3. Txnip was reciprocally increased in CSE^-/-^ mice and decreased by SG1002 treatment. These data suggest that H_2_S positively influences erectile function and penile free radical balance, likely through augmentation of the glutathione and thioredoxin endogenous antioxidant systems.

## 1. INTRODUCTION

Convenience has increasingly become a major driving factor influencing food choices and dietary patterns, which has resulted in heavy consumption of fast, packaged, and processed foods that are rich in added sugar and fat derived from cheap vegetable oils. This dietary pattern is often referred to as the Western diet due to its ascent in the United States and Western Europe, although these dietary trends have spread considerably throughout the globe in recent years.^1^ Pervasiveness of the Western diet is associated with increasing obesity rates and adverse impacts on cardiometabolic health.^2–4^ Epidemiological studies indicate that Western style diets are also associated with increased prevalence of erectile dysfunction (ED).^5,6^ Further, increasing severity of obesity is associated with increasing severity of organic ED, which may be due to a combination of vascular, neurologic, and endocrine deficiencies.^7,8^ As ED often precedes the development of cardiovascular disease, enhanced comprehension of the pathophysiology of ED in response to obesity promoting Western diet conditions may lead to advances in holistic men’s healthcare.

We have previously developed a rodent Western diet (WD) that is reflective of these trends. This diet is high in fat and sugar, with a fatty acid distribution that would be expected from a diet predominantly consisting of fast and processed foods.^9–11^ We have found that both rats and mice develop impaired cavernous nerve-stimulated erectile function within 12 weeks of WD consumption. Excessive production of reactive oxygen species (ROS) is a hallmark of ED pathogenesis.^12^ Indeed, we have observed increased *in vivo* penile ROS in mice following the 12-week WD which was consistent with heightened protein expression of NADPH oxidase (Nox) subunits in the corpus cavernosum.^9^ Further, an acute intracavernous injection of a Nox inhibitor improved nerve-stimulated erectile function in WD-fed rats, implicating a causal role of elevated ROS in erectile function impairment in this model.^11^

Hydrogen sulfide (H_2_S) is an endogenously produced gas that has emerged as an important cell signaling molecule. H_2_S is produced by three enzymes, cystathionine γ-lyase (CSE), cystathionine β-synthase (CBS), and 3-mercaptopyruvate sulfurtransferase (MPST), of which CSE appears to be the predominant source of H_2_S production in the vascular system.^13^ H_2_S has emerged as popular target of investigation in vascular research. While H_2_S exhibits multiple features that may be deemed vasoprotective, CSE expression and associated H_2_S production have been found to be suppressed in several conditions associated with endothelial dysfunction, such as hypertension and diabetes.^13^ Moreover, H_2_S has antioxidant properties that may provide cellular protection against excessive ROS production. H_2_S has been shown to directly scavenge the ROS hydrogen peroxide (H_2_O_2_) and superoxide in an *in vitro* system.^14^ H_2_S may also stimulate transcription of several genes related to ROS detoxification and the cellular antioxidant defense systems.^15^

The advancement of H_2_S-based therapies has faced multiple challenges. First, H_2_S is a gas with a pungent smell characteristic of rotten eggs. Second, H_2_S is a reactive species with a relatively short half-life. Much of the early *in vivo* animal based research focused on H_2_S as a therapeutic has utilized intraperitoneal injections of rapid-release H_2_S donating salts.^16^ In vivo injection of these salts results in large initial spikes of H_2_S, which then decline within minutes.^16,17^ SG1002 is an orally active H_2_S prodrug that releases H_2_S into the bloodstream in a slow and sustained manner. It has additional beneficial aspects in that it is odorless, tasteless, and shelf stable.^18^ Importantly, SG1002 has been shown to be safe and well-tolerated in two clinical trials, one at 800 mg/day for 7 days in heart failure patients, the other at 750 mg/day for 75 days in patients with idiopathic oligoasthenozoospermia.^19,20^ The objectives of this study were to determine 1) if SG1002 treatment reverses WD-associated erectile dysfunction, 2) if SG1002 treatment influences penile ROS levels and the endogenous antioxidant system of the corpus cavernosum, and 3) if CSE deletion influences erectile function, penile ROS levels, and the endogenous antioxidant systems of the corpus cavernosum.

## 2. MATERIALS AND METHODS

### 2.1 Experimental animals and dietary intervention

Male C57Bl/6J mice were purchased from Jackson Laboratories (Bar Harbor, ME, USA). Mice were maintained in a temperature-controlled animal housing facility with ad libitum access to food and drinking water. At 8 weeks of age, mice were randomized into either a Western diet (WD) or a control diet (CD) intervention. The WD (Teklad Diets 110365; Inotiv, Madison, WI, USA) is a high-fat, high-sucrose diet (4.68 kcal/g, 44.6% kcal fat, 40.7% kcal carbohydrate, 340 g/kg sucrose) with a Western pattern fatty acid (39% / 24% / 37% saturated (SFA) / monounsaturated (MUFA) / polyunsaturated (PUFA) fatty acids) distribution enriched with the ω-6 PUFA linoleic acid (27 ω-6:ω-3 ratio). The CD (Teklad Diets 110367) was lower in fat and sucrose (3.67 kcal/g, 12.7% kcal fat, 72.4% kcal carbohydrate, 150 g/kg sucrose) with an approximately equivalent (40% / 26% / 33% SFA/MUFA/PUFA) fatty acid distribution, a moderate 9.6 ω-6:ω-3 ratio, and equivalent levels of vitamins, minerals, and protein when considered on a basis of energy density. The complete composition of these diets has been published previously.^10^ A subgroup of these mice (n=8/group) underwent their respective dietary intervention for 12 weeks, at which point the mice were sacrificed and corpus cavernosum tissue was harvested. The remainder of the mice completed the 12-week dietary intervention, at which point they either remained on their respective diet for an additional 6 weeks or were switched to an equivalent diet with SG1002 (Sulfagenix, Cleveland, OH, USA) added at a concentration designed to deliver approximately 20 mg*kg^-1^*day^-1^ for the subsequent 6 weeks. The study design schematic is illustrated in Figure 1. All mice that underwent the 18-week dietary interventions (n = 35/group) were studied at 26 weeks of age.

**Figure 1.**
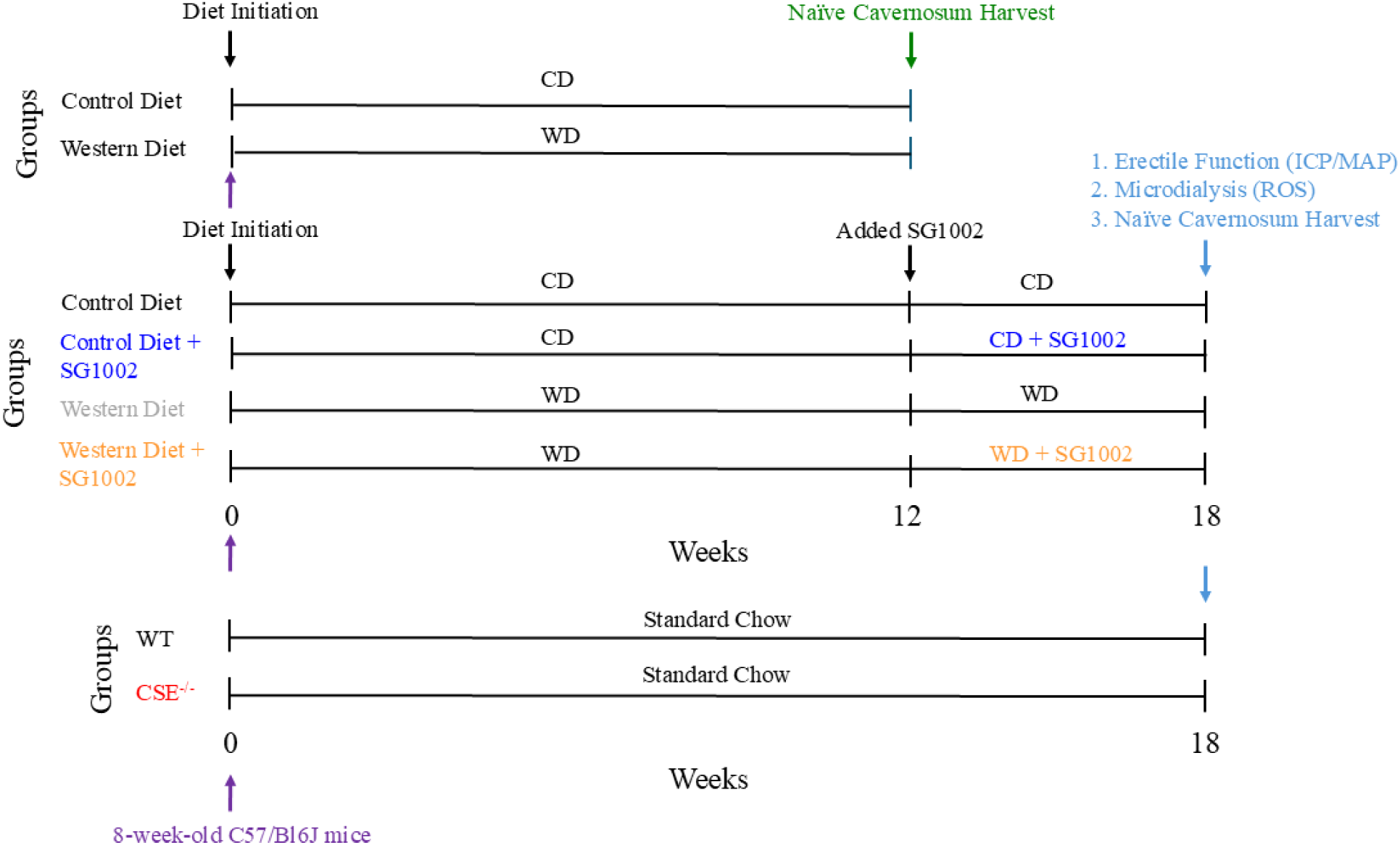
Experimental Design.

In separate studies, CSE knockout (CSE^-/-^) mice were obtained in house using a heterozygous (CSE^+/-^ x CSE^+/-^) breeding scheme, whereby wild type (WT, CSE^+/+^) littermates were used as controls. Mice were maintained with ad libitum access to standard chow diet (5001 PMI Laboratory Rodent Diet, Teklad Diets) and drinking water. Male mice were studied at 26 weeks of age (n = 24/group). Derivation of this strain of mice on the C57/Bl6 background is previously described.^21^ All experimental procedures were approved by the Animal Care and Use Committee of Florida State University.

### 2.2 Metabolic parameters and anthropometrics

Lean mass and fat mass were measured for body composition assessment upon completion of the 18-week dietary intervention in a subset of 15 mice/group by nuclear magnetic resonance-magnetic resonance imaging (EchoMRI 100; Echo Medical Systems, Houston, TX, USA). Blood was obtained from mice in the freely fed state during the euthanasia process via cardiac puncture. Serum concentrations of total cholesterol, triglycerides, high-density lipoprotein cholesterol (HDL-C), low-density lipoprotein cholesterol (LDL-C), very low-density lipoprotein cholesterol (VLDL-C), alanine transaminase (ALT), and aspartate aminotransferase (AST) were obtained with a Piccolo Xpress chemistry analyzer (Abaxis, Union City, CA, USA) using lipid panel plus reagent disks (Abaxis #400-1030) for a subset of 10 mice/group. Serum samples were diluted 1:1 in phosphate buffered saline for analysis.

### 2.3 Erectile function assessment

Erectile function was assessed by measurements of intracavernosal pressure (ICP) and mean arterial pressure (MAP) as described.^22^ Briefly, mice were anesthetized with an intraperitoneal injection of 90 mg/kg ketamine and 10 mg/kg xylazine. The left carotid artery was cannulated with polyethylene tubing filled with 100 U/mL heparinized saline connected to a pressure transducer. The left crus was cannulated with a 30-gauge needle thread into heparinized saline filled polyethylene tubing and connected to a pressure transducer. ICP and MAP were continuously measured with LabChart 8 software (ADInstruments, Sydney, Australia). The prostate was exposed and the right cavernous nerve isolated and gently lifted off of the prostate with platinum bipolar hook electrodes (Natus Neurology, Middleton, WI, USA) using a micromanipulator (World Precision Instruments, Sarasota, FL, USA). The nerve was stimulated with a square pulse stimulator (Grass Instruments, Quincy, MA, USA) for 1 min stimulation periods, separately at 1, 2, and 4 V of stimulation. Erectile function was assessed by the maximal ICP normalized to MAP (ICP/MAP) achieved during each stimulation period.

### 2.4 Penile in vivo ROS measurement

Separate cohorts of mice were used for assessment of *in vivo* ROS production. H_2_O_2_ and superoxide were measured using a microdialysis technique as previously described in detail.^23,24^ Briefly, mice were anesthetized with an intraperitoneal injection of 90 mg/kg ketamine and 10 mg/kg xylazine and placed on a heating pad. The penis was freed of skin and fascia, and a linear microdialysis probe (Bioanalytical Systems, Inc., West Lafayette, IN, USA) with a 3 mm membrane length and 20 kDa maximal pore size was inserted into the shaft of the penis.

Microdialysis probes were perfused with saline containing 100 µM Amplex Ultrared (Molecular Probes, Eugene, OR, USA) and 1.0 U/mL horseradish peroxidase (HRP; Sigma Aldrich, St. Louis, MO, USA) at 1.0 µL/min with a microdialysis pump (CMA Microdialysis, Kista, Sweden). Three 15-min replicate samples were collected, and fluorescence of the dialysate was measured in a fluorometer (BioRad, Hercules, CA, USA) at 510 nm Ex/590 nm Em. Then, 10 U/mL superoxide dismutase (SOD; Sigma Aldrich) was added to the perfusate, which allows for rapid conversion of superoxide that crosses over the microdialysis membrane into H_2_O_2_, which stimulates the conversion of Amplex Ultrared to the fluorescent resorufin.^25,26^ Three 15-min replicate dialysate samples were collected from which fluorescence was measured.

### 2.5 RNA extraction and qRT-PCR

Separate cohorts of mice were used to obtain naïve corpus cavernosum (CC) tissue. Mice were anesthetized with an intraperitoneal injection of 90 mg/kg ketamine and 10 mg/kg xylazine and euthanized by thoracotomy and exsanguination of the vena cava. The penile shaft was exposed by freeing the tissue of skin and fascia, and then isolated by cutting at the penile base and the proximal glans. The corpus spongiosum, dorsal vein, and connective tissues were stripped from the penile shaft. The remaining CC tissue was placed on a Kimwipe, and blood was expelled by gently rolling over the tissue with a curved forceps, rinsed in ice-cold phosphate buffered saline, patted dry, and snap frozen in liquid nitrogen and stored at −80°C until processing. RNA was extracted using a RNeasy mini kit (Qiagen Inc., Hilden, Germany) according to the manufacturer instructions. RNA quantity and purity was determined with a NanoDrop One microvolume spectrophotometer (ThermoFisher Scientific, Waltham, MA, USA). cDNA was synthesized from 300 ng of total RNA using Ready-To-Go You-Prime First Strand Beads (Cytiva, Marlborough, MA, USA). qRT-PCR was conducted on a QuantStudio3 (ThermoFisher Scientific) RT-PCR thermal cycler using TaqMan Fast Advanced Master Mix (ThermoFisher Scientific) as described.^27^ Each sample was performed in triplicate, with relative expression levels of all genes normalized using the ΔΔCt method.^28^ Primer information for all TaqMan gene expression assays are listed in Table S1. Hprt1 was used as the internal control, the expression of which was not affected by WD, SG1002, or CSE knockout.

### 2.6 Immunoblot analysis

Corpus cavernosum tissue samples were homogenized in radioimmunoprecipitation assay buffer (Cell Signaling Technologies, Danvers, MA, USA) with 1 mM phenylmethanesulfonyl fluoride. Homogenates were used for immunoblotting as described.^9^ All antibodies, manufacturer, catalog numbers, and final dilutions used are listed in Table S2.

### 2.7 In vitro amperometric H_2_S measurement

Corpus cavernosum tissue homogenate from the wild type mice that completed the 12-week dietary intervention were used for *in vitro* CSE-mediated H_2_S production measurement using a 3-mm H_2_S-selective electrode connected to an Apollo 4000 Free Radical Analyzer (World Precision Instruments). 160 µg total protein from each sample was brought up to 200 µl total volume with homogenization buffer. Homogenate was incubated with 2 mM pyridoxal-5’-phosphate and 50 mM L-cysteine at 37°C for 50 min in 2-ml gas-tight vials, as described.^29^ Following incubation, 200 µl of 10X PBS adjusted to pH 6.0 were added to the reaction to promote release of H_2_S and to stop the reaction. The reaction was incubated at 37°C for another 10 min. 2 ml of headspace gas was withdrawn from the gas-tight vial using a 10 ml syringe and injected into a scintillation vial containing 15 ml of 10X PBS in which the amperometric probe was equilibrating. H_2_S production was calculated as the difference between the substrate incubated with homogenate and the baseline signal of the substrate with no homogenate. A calibration curve for the amperometric probe was generated using the H_2_S donor sodium hydrosulfide (NaHS) and assuming that one-third of NaHS is in soluble H_2_S form to equate the amperometric signal to an H_2_S concentration.

### 2.8 Data and statistical analysis

Statistical analyses were performed with GraphPad Prism (Prism 10.0, GraphPad, San Diego, CA, USA). For comparisons between two groups, an unpaired t-test was used to determine group differences. In cases where an F test determined significant differences in variance between the two groups, the non-parametric Mann-Whitney U test was used. A two-way repeated measures ANOVA (group x voltage) with Tukey’s multiple comparisons posthoc test was used to determine group differences. For all other measures involving the combination of dietary intervention and SG1002 treatment, a two-way ANOVA (diet x SG1002) was used. Significant main effects and interaction effects are reported. In cases where a significant main effect was present, Sidak’s multiple comparisons posthoc testing was used to determine individual group differences. In all cases, p < 0.05 was considered statistically significant.

## 3. RESULTS

### 3.1 Metabolic Parameters

Final body weight was not different between WT and CSE^-/-^ mice (WT: 34.9±4.3g, CSE^-/-^: 35.9±3.9g; p=0.525). WT mice that completed the 12-week WD intervention gained significant weight compared to their counterparts that remained on the CD (CD: 31.0±1.7g, WD: 44.3±2.3g; p<0.0001). More complete metabolic profiling was performed on mice that completed the 18-week dietary intervention, which is illustrated in Table 1. Final body weight, lean mass, fat mass, and body fat percentage were all elevated following the WD, with no impact of SG1002 observed for these measures. Total cholesterol, HDL-C, LDL-C, and the liver enzymes ALT and AST were all significantly elevated in mice fed the WD without SG1002. These measures were subtly, though not significantly reduced in mice fed the WD with SG1002. There was no effect of SG1002 in the CD condition for any measures of body composition or metabolic profile. Total average daily SG1002 consumed was 19.4±2.0 mg*kg^-1^*day^-1^ when mixed into the CD and 18.9±2.2 mg*kg^-1^*day^-1^ when mixed into the WD.

**Table 1.**
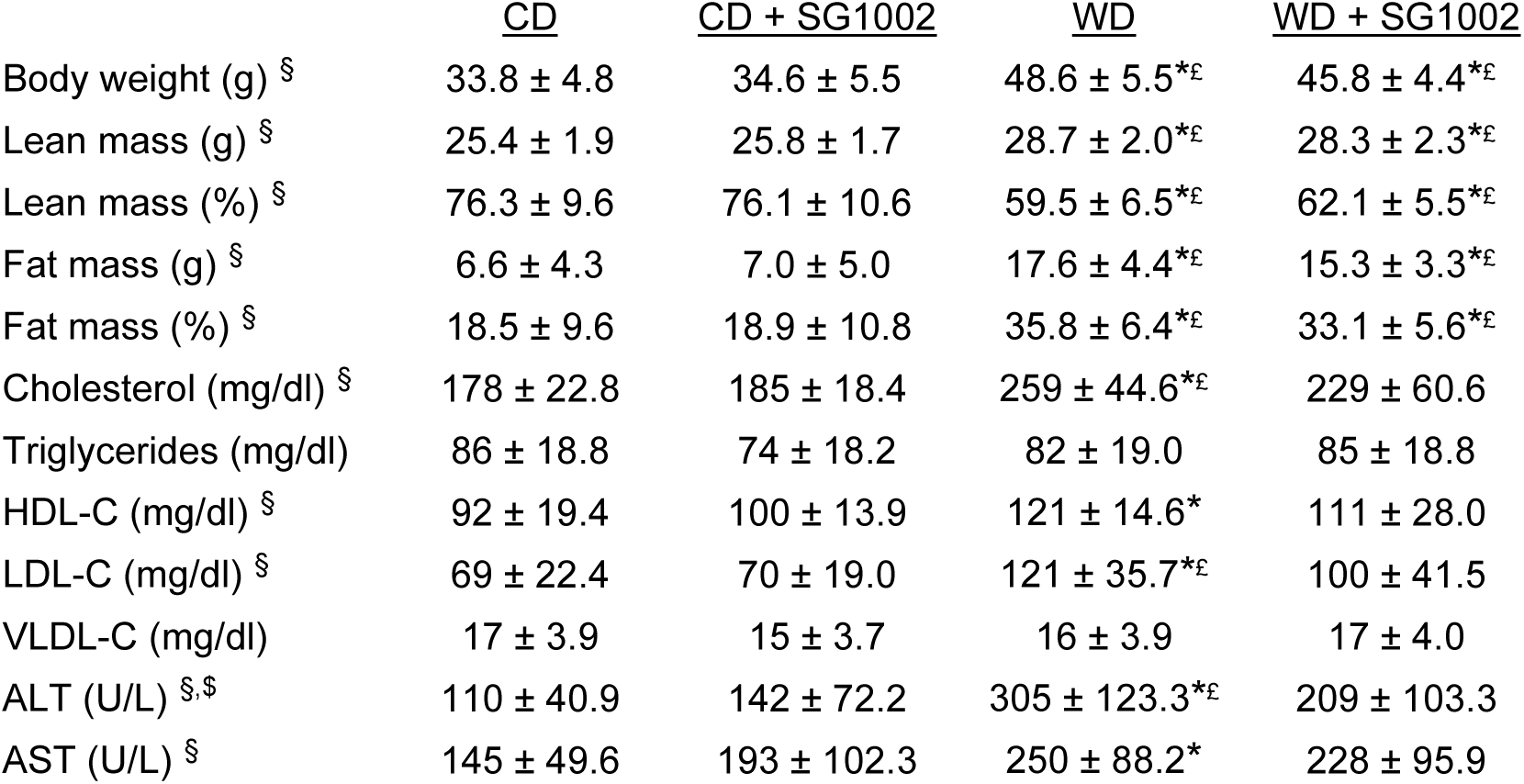
Body composition and serum metabolic profile of mice following the 18-week dietary intervention. Serum was obtained in the freely fed state. Mean ± SD for a subset of n=10 mice/group. ^§^ p<.05 two-way ANOVA main effect of diet, ^$^ p<.05 two-way ANOVA interaction effect. *p<.05 vs. CD. ^£^p<.05 vs. CD+SG1002.

### 3.2 The Western diet diminishes CSE in the corpus cavernosum

We fed mice the control or Western diet for 12 weeks to characterize the protein content of the H_2_S producing enzymes in the CC at this time point consistent with impaired erectile function. Representative immunoblot images are presented in Figure 2A, while the relative quantification of normalized band intensity density is presented in Figure 2B. CSE content was decreased in mice fed the Western diet (p=.006), while no differences were observed for CBS or MPST. Following up on this observation we measured CSE-mediated H_2_S production via amperometry from the CC homogenate (Figure 2C), which was depressed in mice fed the Western diet (p=.032). There were no differences in CBS or MPST gene or protein expression between WT and CSE^-/-^ mice (Figure S3), or between any groups that completed the 18-week dietary intervention (Figure S3). CSE gene (Cth) and protein expression remained depressed following 18-weeks of the WD. SG1002 stimulated an increase in Cth gene and CSE protein expression in the CD condition (Figure S4).

**Figure 2.**
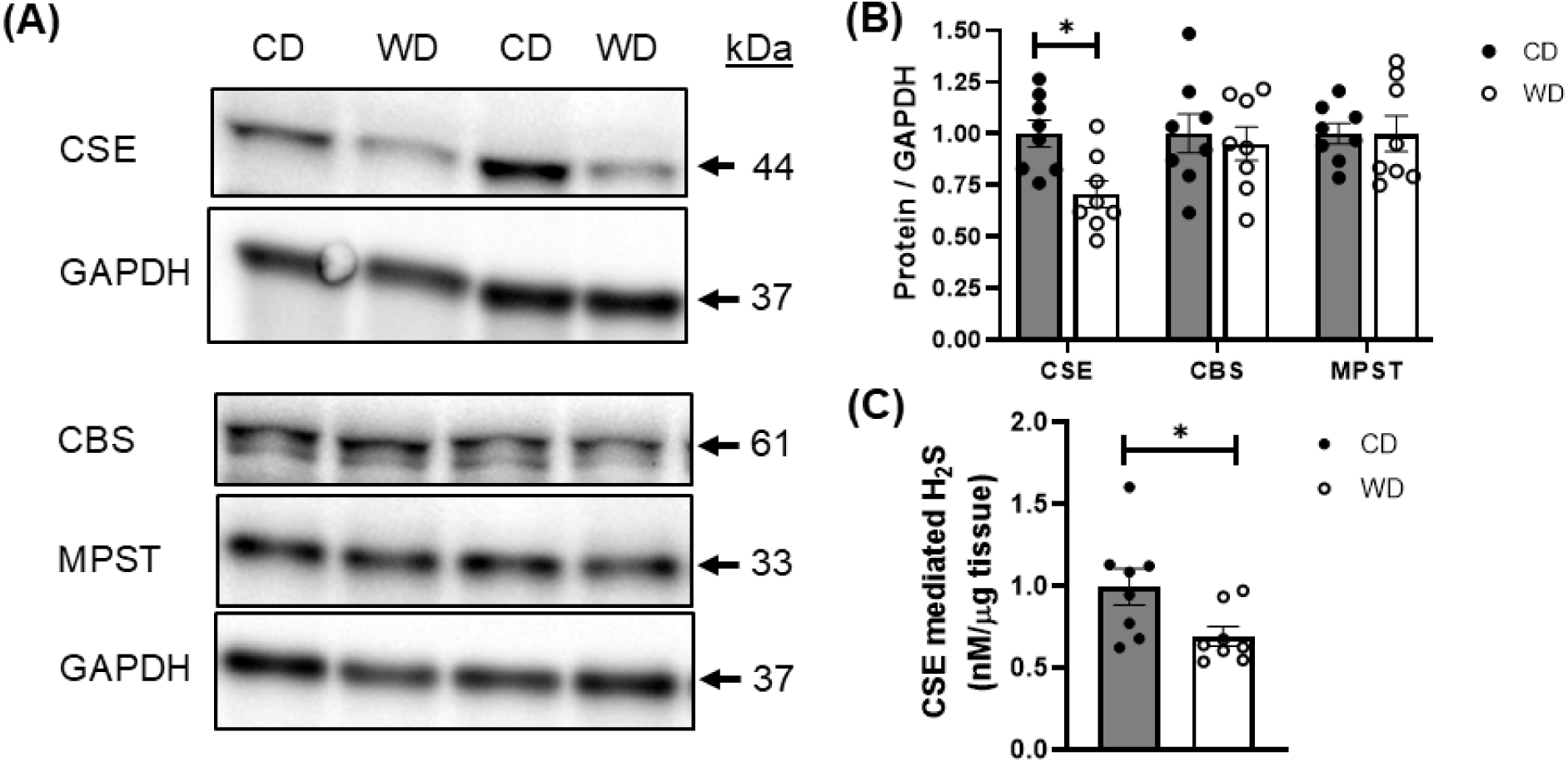
CSE-mediated H_2_S production is diminished in the corpus cavernosum following 12 weeks of the western diet. (A) Representative images of western blots run for the H_2_S production enzymes CSE, CBS, and MPST, as well as GAPDH to serve as the loading control. (B) Quantification of normalized band intensity density. (C) CSE-stimulated H_2_S production was measured from corpus cavernosum homogenate. Bars represent mean ± SEM for n = 8/group. * p<.05 vs. CD.

### 3.3 H_2_S influences erectile function

Representative tracings that include the ICP and MAP readings during cavernous nerve stimulation are presented in Figures S1 and S2. Erectile function was assessed as peak ICP/MAP achieved during nerve stimulation, which was significantly impaired in CSE^-/-^ mice at the medium (2V) and high (4V) doses of stimulation (Figure 3A). Erectile function was impaired throughout the voltage range in wild type mice fed the Western diet for 18 weeks (Figure 3B). Erectile function was improved throughout the voltage range in Western diet fed mice treated with SG1002 for the final 6 weeks. These improvements represent 86% (1V), 53% (2V), and 51% (4V) mean restorations of the CD values. There were no significant effects of SG1002 treatment in mice fed the control diet.

**Figure 3.**
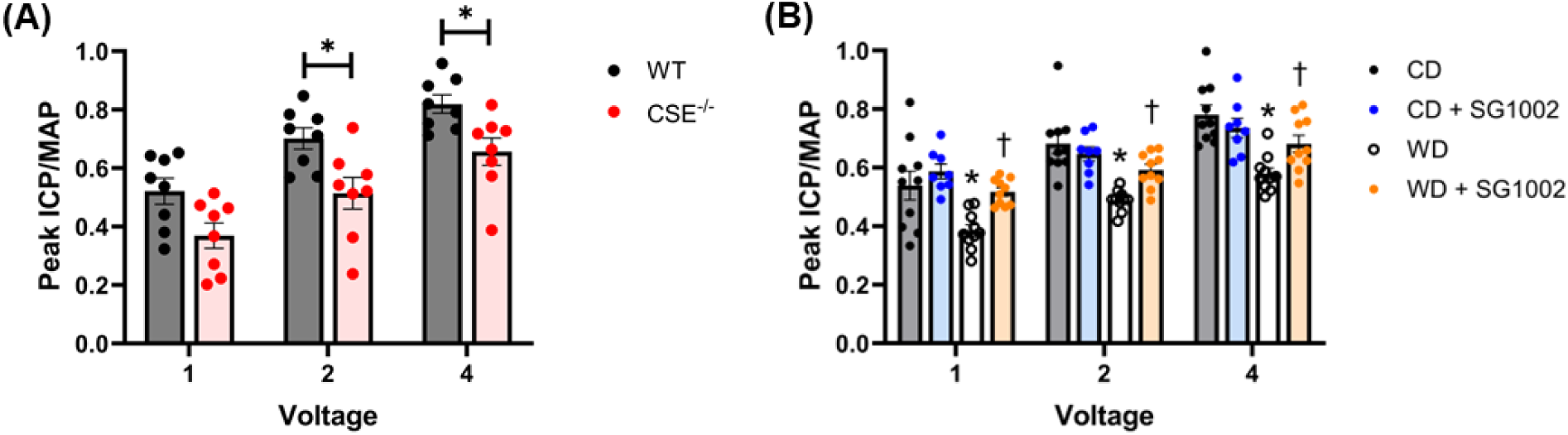
Influence of H_2_S on erectile function. Erectile function was assessed by electrical stimulation of the cavernous nerve at 1, 2, and 4 volts, and expressed as the peak intracavernous pressure (ICP) normalized to mean arterial pressure (MAP). (A) Erectile function in wild type (WT) mice and mice deficient in cystathionine γ-lyase (CSE^-/-^), (n = 8/group). * p<.05 vs. WT. (B) Erectile function following the 18-week control diet (CD) or Western diet (WD) with and without the H_2_S prodrug SG1002 for the final 6 weeks. Bars represent mean ± SEM for n = 8-10/group. * p<.05 CD vs. WD. ^†^ p<.05 WD vs. WD + SG1002.

### 3.4 H_2_S suppresses penile ROS levels

There was a trend (p=.051) toward elevated *in vivo* penile H_2_O_2_ levels in CSE^-/-^ mice relative to WT littermate control mice (Figure 4A). Addition of superoxide dismutase (SOD) to the microdialysis perfusate allows for additional contribution of superoxide that crosses into the microdialysis membrane to the total ROS fluorescent signal. Under these conditions, the total penile ROS signal captured was increased in CSE^-/-^ mice (p=.001, Figure 4B). The increase in signal upon SOD addition is attributed to superoxide (Figure 4C), which was also elevated CSE^-/-^ mice (p=.020). Data for *in vivo* penile H_2_O_2_, total ROS, and superoxide for mice that underwent the 18-week dietary intervention are presented in Figure 4D-F. Two-way ANOVA revealed a main effect of diet for all three of these measures, indicating a propensity toward increased penile ROS induced by the Western diet. There was also a main effect of SG1002 treatment observed for all measures, indicating a suppressive effect of SG1002 on penile ROS levels. Post-hoc analysis revealed an increase in all of these measures in the WD group compared to the CD group, while SG1002 treatment in the Western diet condition blunted all of these values (WD+SG1002 vs. WD). Post-hoc analysis further revealed that the total ROS measurement was increased in WD-fed mice treated with SG1002 compared to CD-fed mice treated with SG1002.

**Figure 4.**
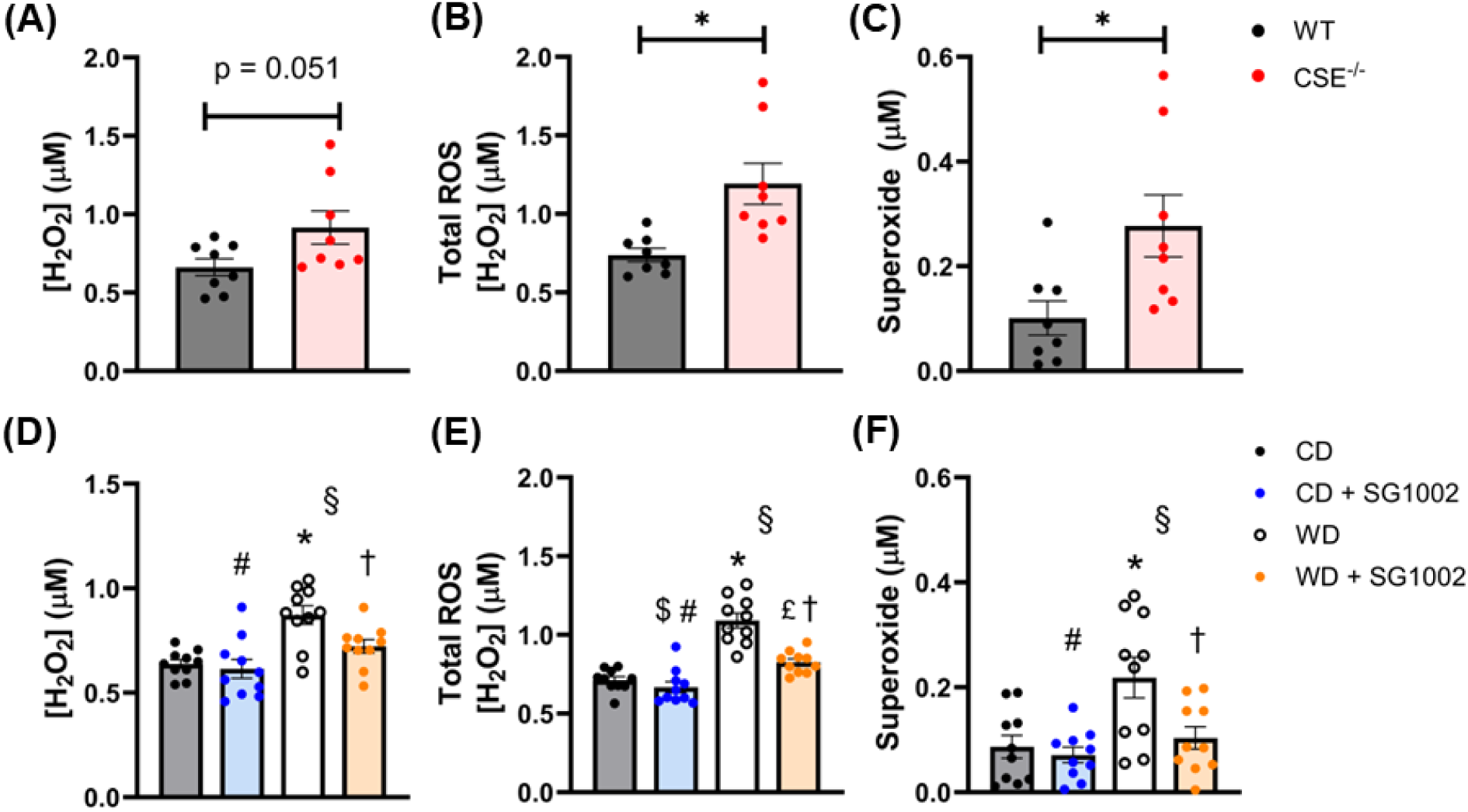
Influence of H_2_S on penile reactive oxygen species (ROS) levels. Penile levels of the ROS hydrogen peroxide (H_2_O_2_) and superoxide (O_2_^•-^) were measured *in vivo* using a microdialysis technique coupled to the Amplex Ultrared assay with and without superoxide dismutase. Measures taken in wild type (WT) mice and mice deficient in cystathionine γ-lyase (CSE^-/-^), (n = 8/group) for (A) H_2_O_2_, (B) Total ROS, and (C) superoxide. * p<.05 vs. WT. Measures taken following the 18-week control diet (CD) or Western diet (WD) with and without the H_2_S prodrug SG1002 for the final 6 weeks for (D) H_2_O_2_, (E) Total ROS, and (F) superoxide. Bars represent mean ± SEM for n = 10/group. # p<.05 two-way ANOVA main effect of SG1002 treatment, § p<.05 two-way ANOVA main effect of diet, $ p<.05 two-way ANOVA interaction effect. The following indicate a significant difference between groups via post-hoc analysis: * CD vs. WD; ^†^ WD vs. WD + SG1002; £ CD + SG1002 vs. WD + SG1002.

### 3.5 H_2_S stimulates cellular antioxidant defense in the corpus cavernosum

Representative images and relative protein quantification from immunoblotting experiments, as well as expression of genes related to cellular antioxidative cytoprotection are presented in Figure 5A-C (WT vs. CSE^-/-^) and Figure 6A-C (18-week dietary intervention). We observed significant decreases in gene expression of NADPH quinone dehydrogenase 1 (Nqo1), glutamate-cysteine ligase catalytic subunit (Gclc), glutathione S-transferase mu-1 (Gstm1), glutathione peroxidase 1 (Gpx1), glutathione peroxidase 4 (Gpx4), and heme oxygenase-1 (Hmox1) in the CC of CSE^-/-^ mice relative to WT littermate controls (Figure 5B). There was no difference observed for glutamate-cysteine ligase modifier subunit (Gclm). We were able to successfully attain immunoblots for GCLC, GPX4, and heme oxygenase-1 (protein HO-1), which verified that these differences remained at the protein level (Figure 5C). When investigating these cytoprotective antioxidant genes in the wild type mice following the 18-week dietary intervention, we observed a two-way ANOVA main effect of diet for Nqo1 gene expression, indicating that the Western diet reduced Nqo1 levels (Figure 6B). There were no other significant effects of the diet. We observed a two-way ANOVA main effect of SG1002 treatment for Nqo1, Gclc, Gstm1, Gpx1, Gpx4, and Hmox1, whereby SG1002 stimulated an increase in expression of all of these genes. This SG1002 stimulatory effect remained at the protein level for the proteins investigated (GCLC, GPX4, HO-1; Figure 6C).

**Figure 5.**
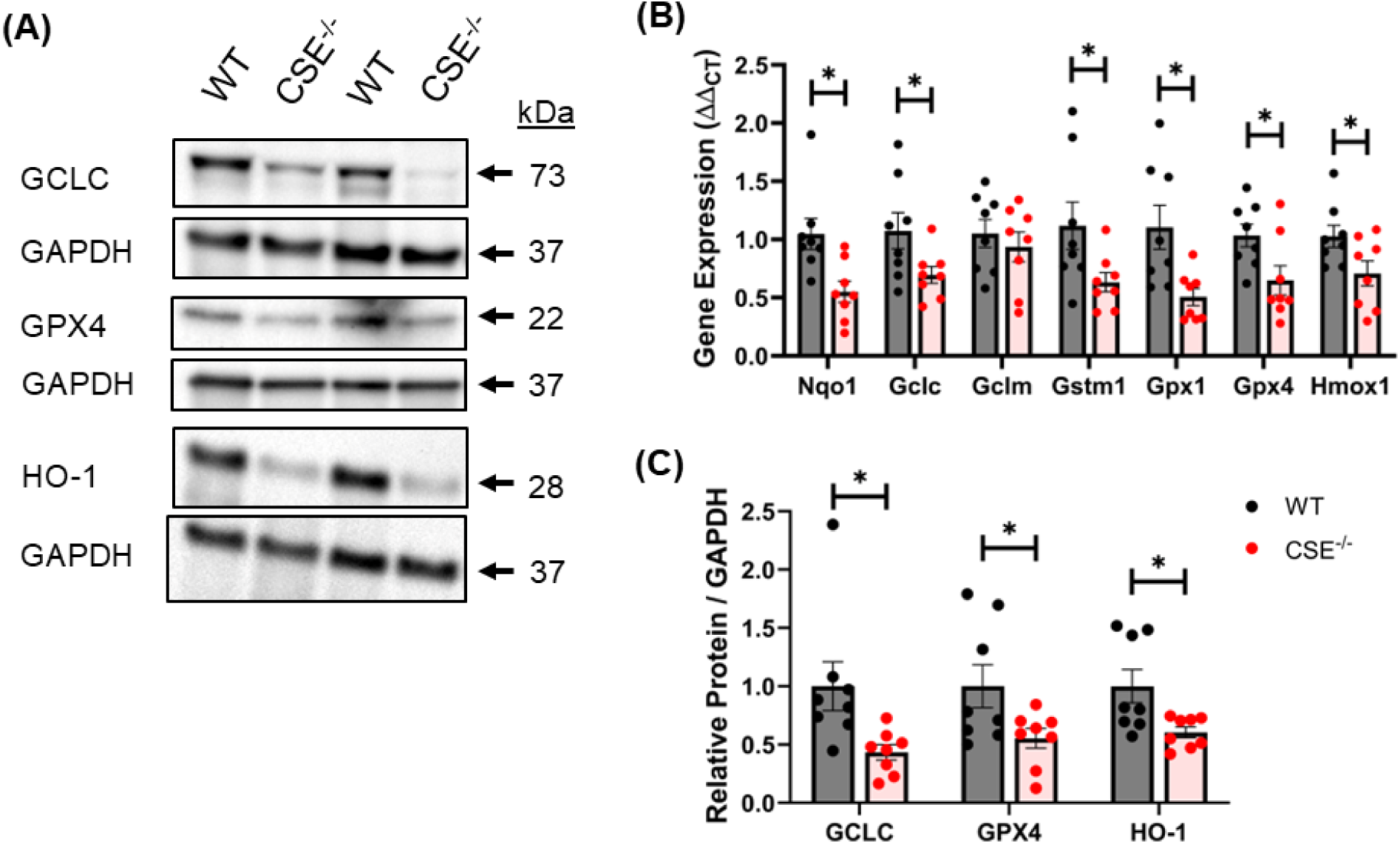
Cystathionine γ-lyase (CSE) deficiency diminishes cellular cytoprotective antioxidant gene and protein expression in the corpus cavernosum. (A) Representative images of western blots run for GCLC, GPX4, and HO-1, as well as the respective loading control protein GAPDH. (B) Expression of the cytoprotective genes Nqo1, Gclc, Gclm, Gstm1, Gpx1, Gpx4, and Hmox1 assessed by qRT-PCR. (C) Quantification of normalized band intensity density of proteins assessed by western blot. Bars represent mean ± SEM for n = 8/group. * p<.05 vs. WT.

**Figure 6.**
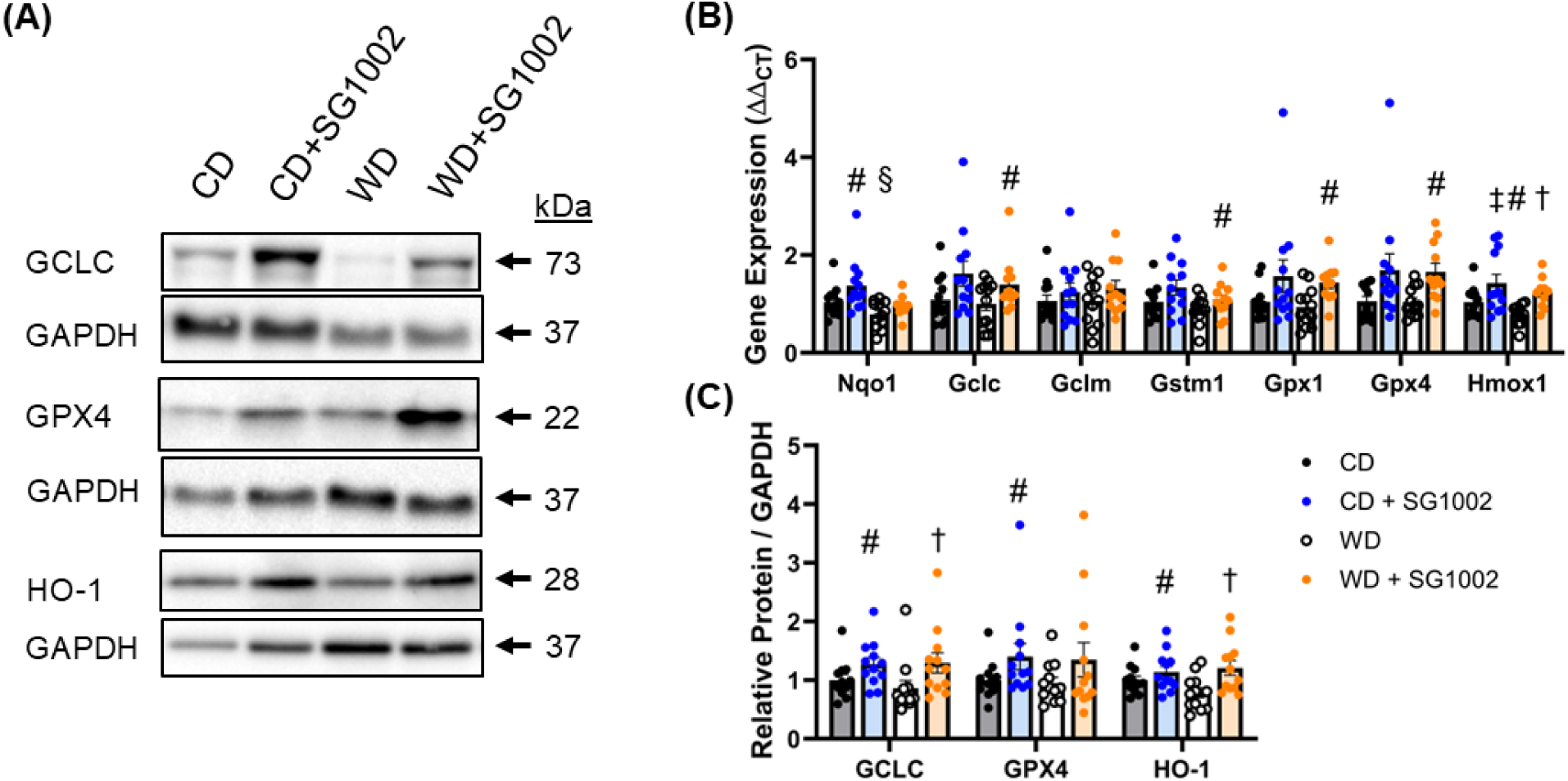
H_2_S therapy augments cellular cytoprotective antioxidant gene and protein expression in the corpus cavernosum. Measures taken following the 18-week control diet (CD) or Western diet (WD) with and without the H_2_S prodrug SG1002 for the final 6 weeks. (A) Representative images of western blots run for GCLC, GPX4, and HO-1, as well as the respective loading control protein GAPDH. (B) Expression of the cytoprotective genes Nqo1, Gclc, Gclm, Gstm1, Gpx1, Gpx4, and Hmox1 assessed by qRT-PCR. (C) Quantification of normalized band intensity density of proteins assessed by western blot. Bars represent mean ± SEM for n = 11-12/group. # p<.05 two-way ANOVA main effect of SG1002 treatment, § p<.05 two-way ANOVA main effect of diet. The following indicate a significant difference between groups via post-hoc analysis: ^†^ WD vs. WD + SG1002; ^‡^ CD vs. CD + SG1002.

Representative images and relative protein quantification from immunoblotting experiments, as well as expression of genes related to oxidant dismutation and antioxidant gene regulation are presented in Figure 7A-C (WT vs. CSE^-/-^) and Figure 8A-C (18-week dietary intervention). No differences were observed for gene expression of catalase (Cat), superoxide dismutase 2 (Sod2), superoxide dismutase 3 (Sod3), sirtuin 1 (Sirt1), or sirtuin 3 (Sirt3) in the CC of CSE^-/-^ mice compared to WT littermate controls (Figure 7B). Immunoblotting similarly indicated no differences in SOD2, SOD3, SIRT1, or SIRT3 at the protein level. We did observe a significant decrease in gene expression of superoxide dismutase 1 (Sod1) in CSE^-/-^ mice, however this difference did not hold at the protein level (p=.107). When investigating these gene and protein levels in the wild type mice following the 18-week dietary intervention, we did not observe any significant effects of the Western diet for any of these genes or proteins. We observed a two-way ANOVA main effect of SG1002 treatment for Sirt3 gene expression, however this effect did not translate to the protein level. No other significant effects of SG1002 treatment were observed for gene or protein expression.

**Figure 7.**
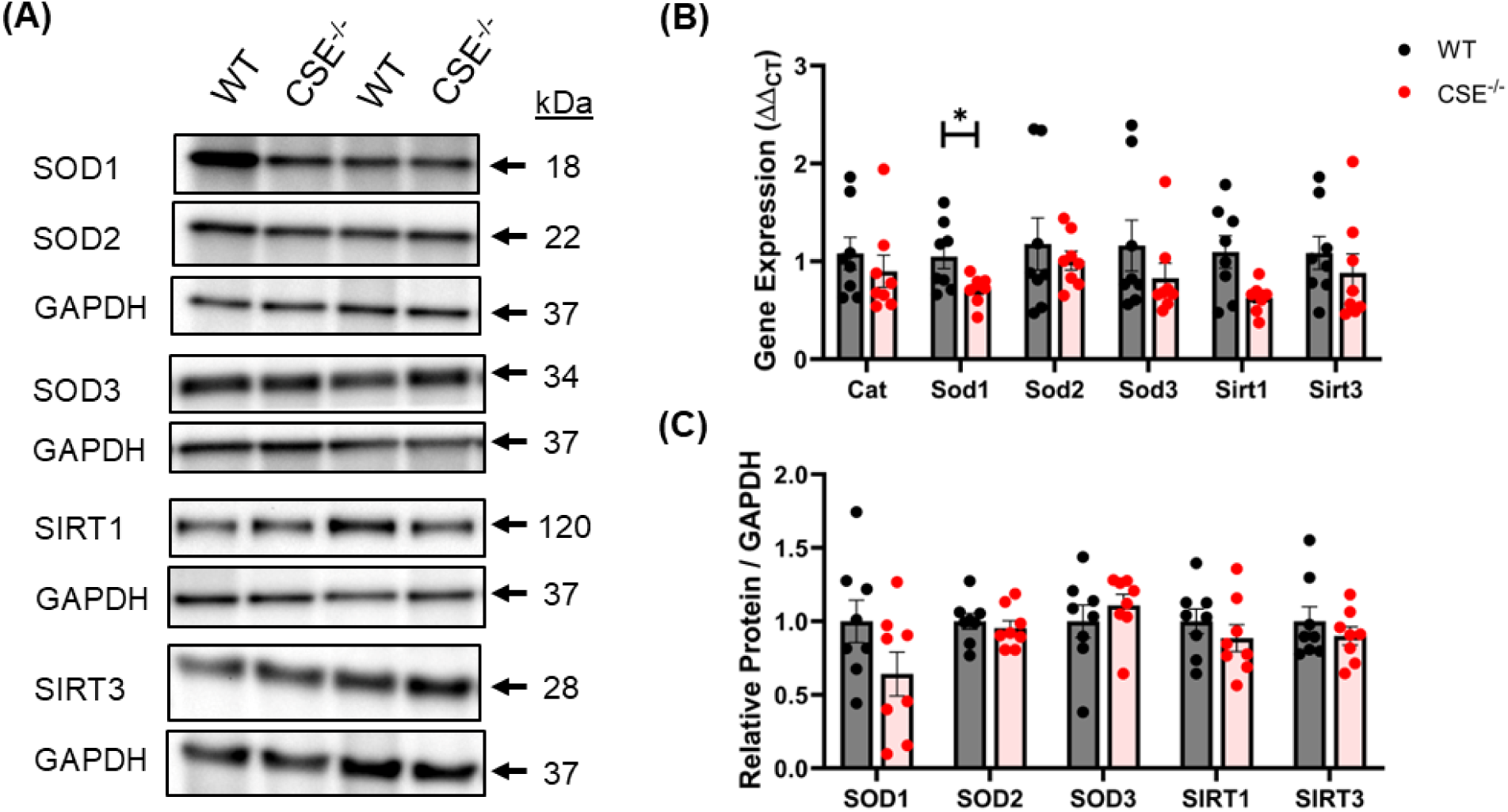
Cystathionine γ-lyase (CSE) deficiency marginally impacts oxidant dismutation and antioxidant regulatory gene and protein expression in the corpus cavernosum. (A) Representative images of western blots run for SOD1, SOD2, SOD3, SIRT1, and SIRT3, as well as the respective loading control protein GAPDH. (B) Gene expression of Cat, Sod1, Sod2, Sod3, Sirt1, and Sirt3 assessed by qRT-PCR. (C) Quantification of normalized band intensity density of proteins assessed by western blot. Bars represent mean ± SEM for n = 8/group. * p<.05 vs. WT.

**Figure 8.**
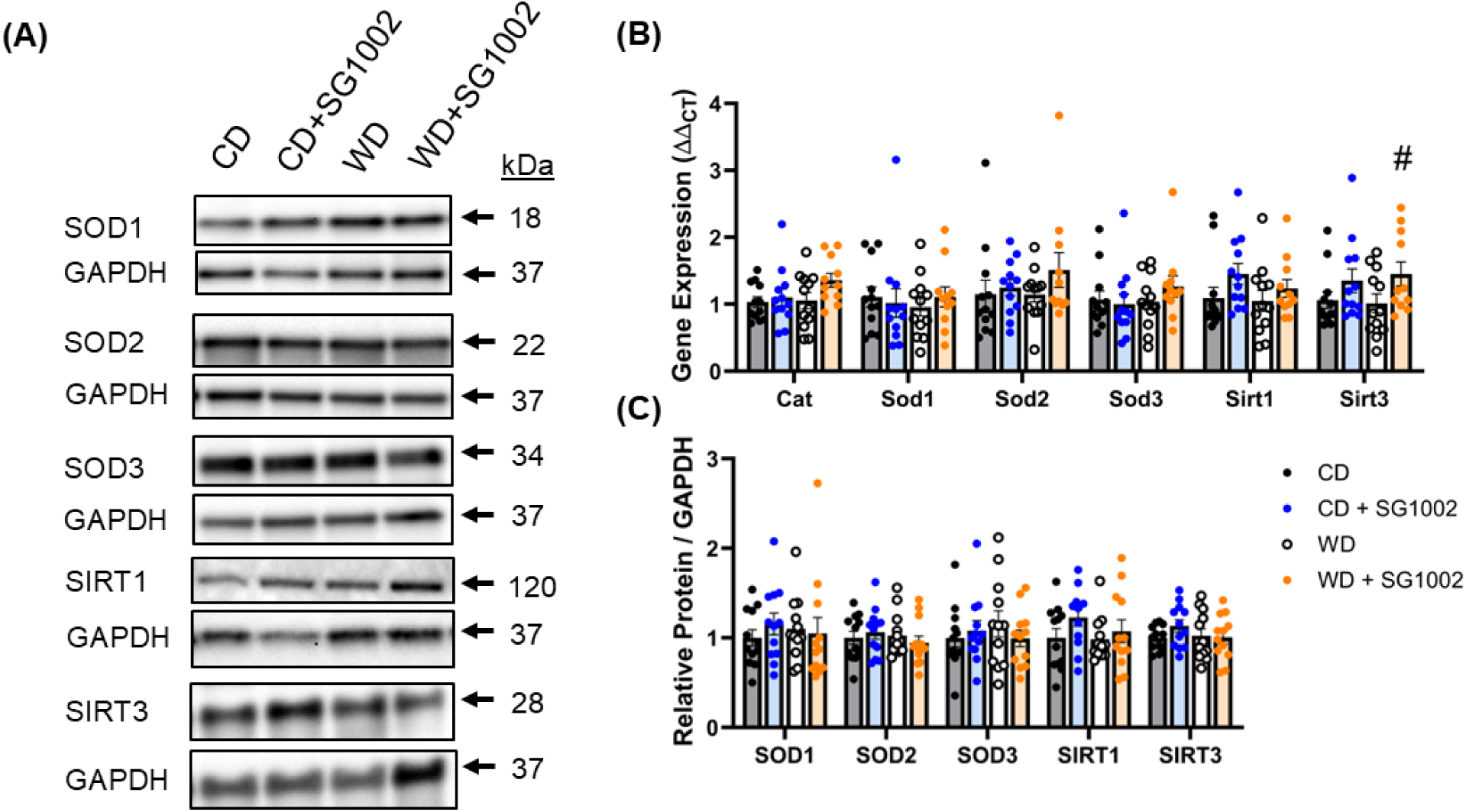
H_2_S therapy marginally impacts oxidant dismutation and antioxidant regulatory gene and protein expression in the corpus cavernosum. Measures taken following the 18-week control diet (CD) or Western diet (WD) with and without the H_2_S prodrug SG1002 for the final 6 weeks. (A) Representative images of western blots run for SOD1, SOD2, SOD3, SIRT1, and SIRT3, as well as the respective loading control protein GAPDH. (B) Gene expression of Cat, Sod1, Sod2, Sod3, Sirt1, and Sirt3 assessed by qRT-PCR. (C) Quantification of normalized band intensity density of proteins assessed by western blot. Bars represent mean ± SEM for n = 11-12/group. # p<.05 two-way ANOVA main effect of SG1002 treatment.

Representative images and relative protein quantification from immunoblotting experiments, as well as expression of genes related to the thioredoxin antioxidant system are presented in Figure 9A-C (WT vs. CSE^-/-^) and Figure 10A-C (18-week dietary intervention). We observed significant decreases in thioredoxin reductase 1 (Txnrd1), thioredoxin reductase 2 (Txnrd2), peroxiredoxin 3 (Prdx3), and peroxiredoxin 5 (Prdx5), while we observed an increase in thioredoxin-interacting protein (Txnip) in the CC of CSE^-/-^ mice compared to WT littermate controls (Figure 9B). No differences in gene expression were observed for thioredoxin 1 (Txn1) or thioredoxin 2 (Txn2). The increase in TXNIP and the decreases in PRDX3 and PRDX5 in CSE^-/-^ mice were also observed at the protein level, although TXNRD1 protein was not different between groups (Figure 9C). When investigating these genes of the thioredoxin antioxidant system in the wild type mice following the 18-week dietary intervention, we observed a two-way ANOVA main effect of SG1002 treatment for Txnrd1, Prdx3, and Txnip, whereby SG1002 stimulated expression of Txnrd1 and Prdx3, and suppressed expression of Txnip (Figure 10B). We observed a two-way ANOVA interaction effect on Txnrd2 gene expression. We also observed a two-way ANOVA main effect of diet on Prdx3 gene expression, whereby the Western diet stimulated Prdx3 gene expression. There were no significant effects related to Txn1, Txn2, or Prdx5 gene expression. The stimulatory two-way ANOVA main effect of SG1002 treatment for TXNRD1 and PRDX3 was also observed at the protein level (Figure 10C). The suppressive effect of SG1002 treatment on TXNIP was also observed at the protein level. It should be noted that post-hoc analysis also revealed increases in TXNIP in the WD group compared to the CD group, as well as a blunting of TXNIP in the WD+SG1002 group compared to the WD group at the gene and protein level.

**Figure 9.**
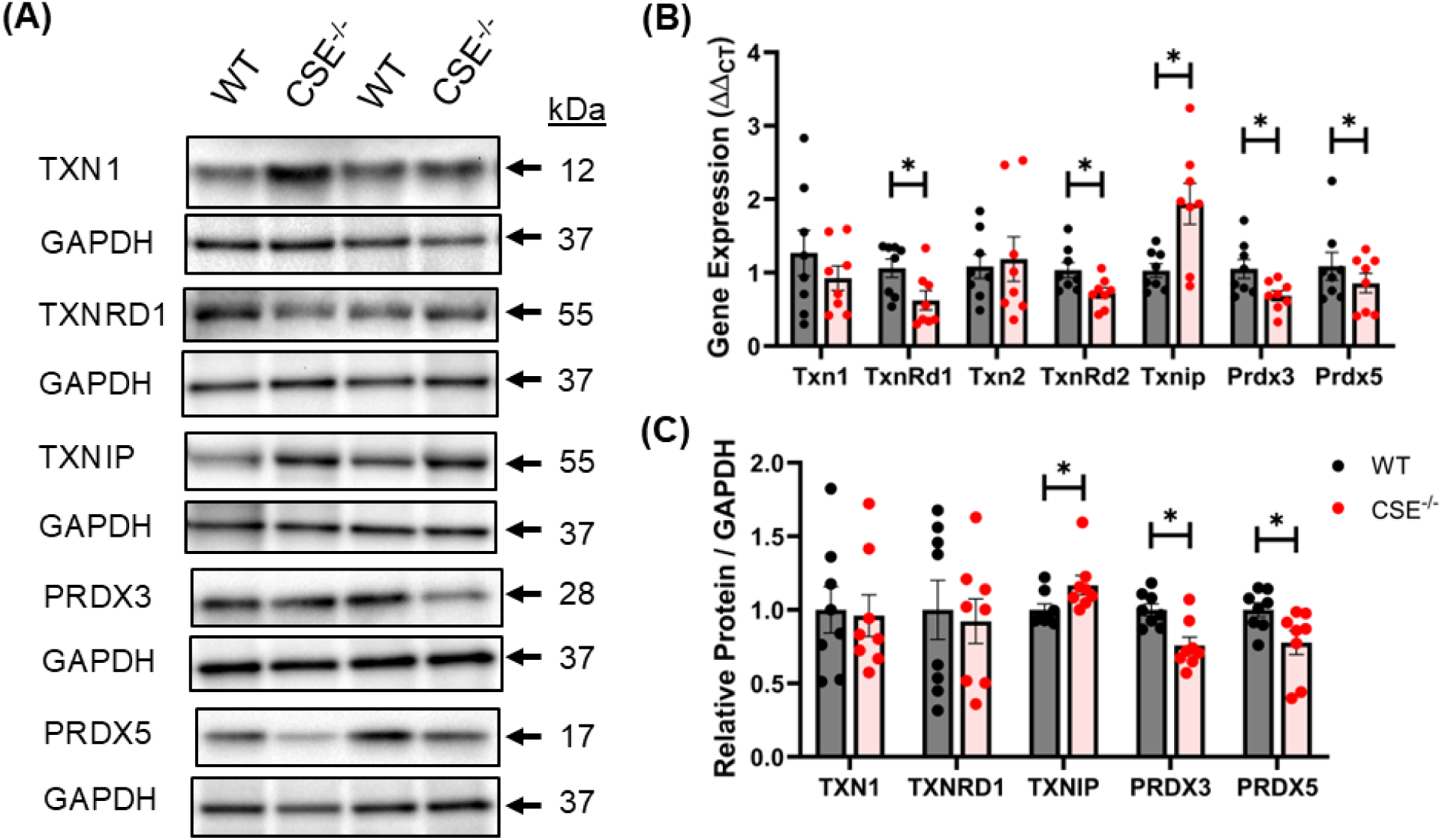
Cystathionine γ-lyase (CSE) deficiency alters gene and protein expression of the thioredoxin antioxidant system in the corpus cavernosum. (A) Representative images of western blots run for TXN1, TXNRD1, TXNIP, PRDX3, and PRDX5, as well as the respective loading control protein GAPDH. (B) Gene expression of Txn1, TxnRd1, Txn2, TxnRd2, Txnip, Prdx3, and Prdx5 assessed by qRT-PCR. (C) Quantification of normalized band intensity density of proteins assessed by western blot. Bars represent mean ± SEM for n = 8/group. * p<.05 vs. WT.

**Figure 10.**
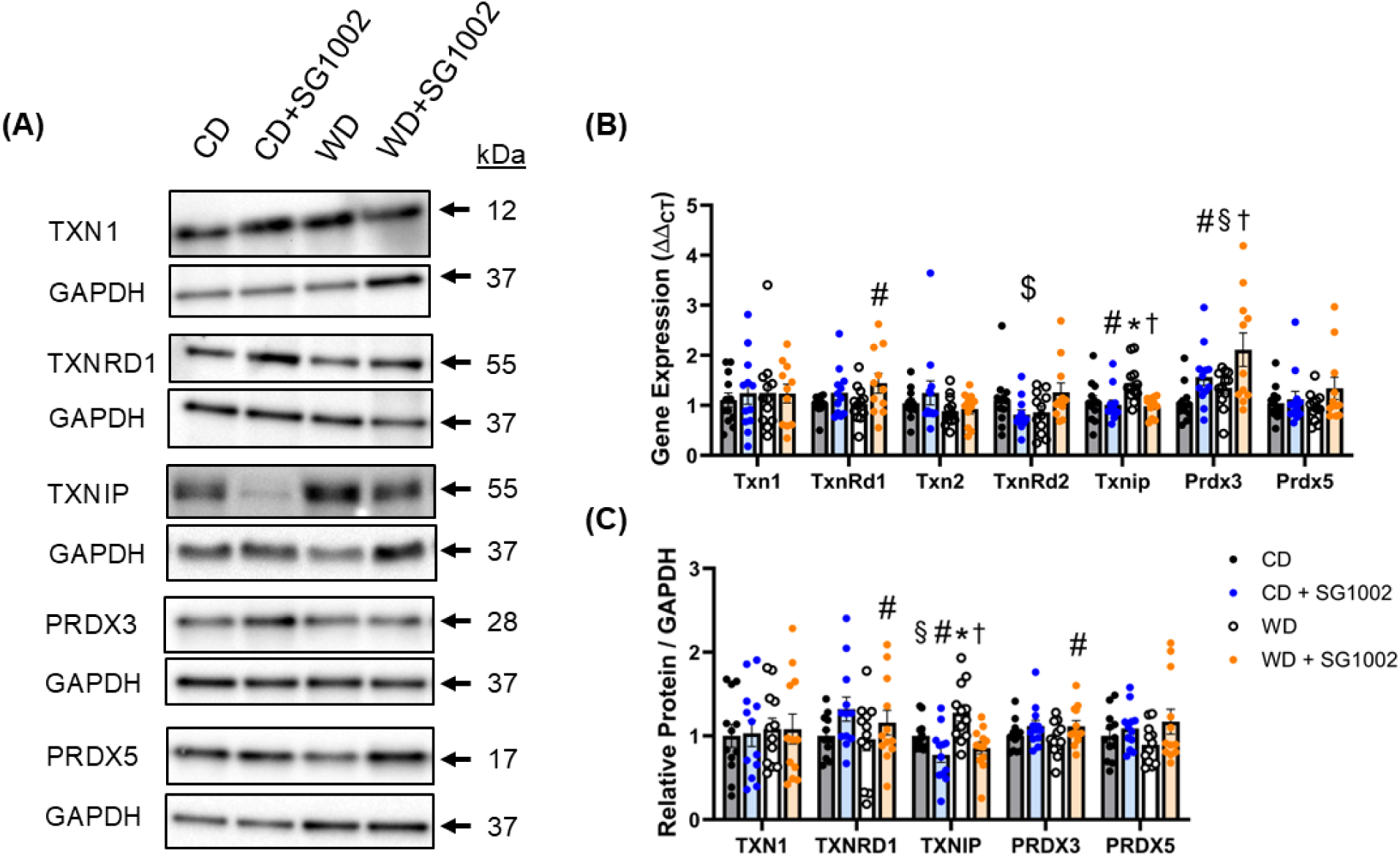
H_2_S therapy alters gene and protein expression of the thioredoxin antioxidant system in the corpus cavernosum. Measures taken following the 18-week control diet (CD) or Western diet (WD) with and without the H_2_S prodrug SG1002 for the final 6 weeks. (A) Representative images of western blots run for TXN1, TXNRD1, TXNIP, PRDX3, and PRDX5, as well as the respective loading control protein GAPDH. (B) Gene expression of Txn1, TxnRd1, Txn2, TxnRd2, Txnip, Prdx3, and Prdx5 assessed by qRT-PCR. (C) Quantification of normalized band intensity density of proteins assessed by western blot. Bars represent mean ± SEM for n = 11-12/group. # p<.05 two-way ANOVA main effect of SG1002 treatment, § p<.05 two-way ANOVA main effect of diet, $ p<.05 two-way ANOVA interaction effect. The following indicate a significant difference between groups via post-hoc analysis: * CD vs. WD; ^†^ WD vs. WD + SG1002.

### 3.6 H_2_S marginally influences expression of oxidant proteins in the corpus cavernosum

Representative images and relative protein quantification from immunoblotting experiments for xanthine oxidase and NADPH oxidase subunits are presented in Figure S5A-C (WT vs. CSE^-/-^) and Figure S6A-C (18-week dietary intervention). No differences were observed for any of these complexes between WT and CSE^-/-^ mice. We observed a two-way ANOVA main effect of diet, whereby the Western diet promoted increased expression of Nox2, p22^phox^, and p47^phox^ in the corpus cavernosum. Post-hoc analysis revealed an increased expression of all three of these complexes in the WD group compared to the CD group, as well as an increase in p47^phox^ in the WD+SG1002 group compared to the CD+SG1002 group. There were no significant effects of SG1002 treatment on any of these complexes, although post-hoc analysis revealed a trend toward a decrease in p22^phox^ expression in the WD+SG1002 group compared to the WD group (p=.0502).

## 4. DISCUSSION

In the present study, we investigated the influence of H_2_S on erectile function and penile ROS, a major driver of ED in conditions associated with the metabolic syndrome. We observed a decrement in protein content and activity of CSE, a major enzymatic source of endogenous H_2_S, in the corpus cavernosum of mice following 12 weeks of a westernized diet high in sucrose and fat derived from saturated and ω-6 polyunsaturated fatty acids. Mice with a deletion of CSE demonstrated impaired erectile function and increased penile ROS levels, which was congruent with decreased corpus cavernosum expression of antioxidant genes and proteins, particularly those involved in the function and restoration of the glutathione and thioredoxin antioxidant systems. Treatment of wild type mice following 12 weeks of the western diet with an H_2_S prodrug (SG1002) for the subsequent six weeks partially restored erectile function and suppressed penile ROS levels. SG1002 treatment had a stimulatory effect on corpus cavernosum expression of antioxidant genes and proteins related to the function and restoration of the glutathione and thioredoxin antioxidant systems. We have consistently observed impaired erectile function following 12 weeks of the WD, thus the effects of SG1002 administered at this time point reflect a reversal of impairment, rather than a mere prevention of dysfunction.

Obesity and increasing number of components of the metabolic syndrome are independent risk factors for ED.^30^ In these conditions, arterial inflow insufficiency is a primary cause of ED that is largely attributed to endothelial dysfunction. First line therapies for ED include oral phosphodiesterase type 5 (PDE5) inhibitors which are ineffective in 30-50% of patients, while efficacy of these drugs progressively declines with increasing ED severity and with decreasing cardiometabolic health, thus more efficacious therapeutic options are needed.^31–33^ La Fuente et al. previously observed decreased CSE protein expression in penile tissue of men with severe ED relative to organ donor controls, implicating H_2_S as a potential pharmacologic target for ED treatment.^34^ While SG1002 has not previously been investigated for ED, treatment of a mouse model of H_2_S deficiency with SG1002 has been shown to restore mesenteric artery flow-induced dilation, indicating a therapeutic effect of SG1002 on endothelial function.^35^ Further, 20 mg*kg^-1^*day^-1^ SG1002 treatment improved ejection fraction and left ventricular end systolic diameter in high-fat diet and pressure overload mouse models of heart failure.^36,37^ This was the same dose of SG1002 that was used in the present study, both of which demonstrated reduced circulating and heart tissue contents of free sulfide and sulfane sulfur levels in the heart failure condition which were augmented by SG1002 treatment. Of further relevance, the high-fat diet study used a 24-week intervention and two SG1002 treatment groups, one that received SG1002 throughout the duration of the dietary intervention, and one in which SG1002 was initiated 12 weeks into the intervention, while temporal measures of cardiac performance were taken every four weeks.^36^ Four weeks after the delayed SG1002 treatment there were no significant differences between SG1002 treatment groups, nor were there differences in terminal cardiac function measures taken after the 24-week dietary intervention between the SG1002 treatment groups.^36^ Similar to the present study, SG1002 treatment reversed a physiological impairment in a relatively short period of time. These findings combined with the impaired erectile function in CSE^-/-^ mice implicate an important role for H_2_S in the preservation of erectile health.

Increased ROS and the associated oxidative stress is a hallmark of ED of various etiologies.^38–41^ We measured penile H_2_O_2_ and superoxide in anesthetized mice using microdialysis. This technique samples from the interstitium, therefore the ROS signal attained is not meant to reflect the total intracellular ROS levels, but rather is reflective of the ROS capable of cell-cell signal transduction.^23^ We observed a 62% increase in total ROS detected in the CSE^-/-^ mice relative to the WT mice. While the majority of the signal attained with this technique derives from H_2_O_2_, we observed a greater proportional increase in the superoxide contribution to the total signal in the CSE^-/-^ condition. Prior work has demonstrated increased superoxide levels in cardiac atria tissue of CSE^-/-^ mice,^42^ as well as increases in general markers of oxidative stress in cardiac tissue,^43^ skeletal muscle,^44^ and mesenteric artery smooth muscle cells.^45^ The present study extends these findings into the penile tissue interstitium. Furthermore, we observed a 53% increase in total ROS in mice fed the WD relative to the CD, while SG1002 treatment of WD mice suppressed this increase 70% of the way back to the CD level. The relative contributions of H_2_O_2_ and superoxide to the total ROS signal were both elevated in WD, while there were no discernable differences in how SG1002 affected these individual components. Our prior work in this WD mouse-model suggests that the excessive ROS associated with the WD is predominantly Nox-mediated.^9^ While we did not use a Nox inhibitor in the microdialysis experiments in the present study, we again observed an increase in cavernous expression of the Nox2 protein, as well as the Nox2 subunits p22^phox^ and p47^phox^ following the WD. None of the oxidant proteins investigated were impacted by CSE^-/-^, while the only notable impact of SG1002 treatment was a trending (p=.0502) decrease in p22^phox^ in the WD condition. Prior work investigating the effects of various H_2_S donors on these Nox proteins has predominantly focused on Nox2. H_2_S donors have been found to decrease Nox2 expression in the renal artery of a hypertension model and myocardium in the streptozotocin type 1 diabetic model, although a fast-releasing H_2_S donor had no impact on Nox2 expression in the streptozotocin model aorta.^46–48^ Treatment with the H_2_S donor erucin suppressed p22^phox^ expression in endothelial cells exposed to high glucose conditions, demonstrating a precedence for an effect of H_2_S on p22^phox^.

While the impact of CSE deficiency and H_2_S therapy on potential sources of ROS in the cavernosum appeared to be marginal, the more likely influence on penile ROS balance is through alteration of cellular antioxidant defense in the cavernosum. We observed several instances whereby antioxidant protein and/or gene expressions were decreased in the CSE^-/-^ condition, while many of these were augmented with SG1002 treatment regardless of dietary condition. Collectively, our results point toward a stimulatory effect of H_2_S on the ability of glutathione to detoxify ROS in the corpus cavernosum. Gclc and Gclm form glutamate-cysteine ligase, an enzyme that catalyzes the rate limiting step in glutathione biosynthesis. While we observed no effect of H_2_S on the modifier subunit (Gclm), a decrement in the catalytic subunit (Gclc), as observed in the CSE^-/-^ cavernosum, may result in impaired glutathione biosynthesis. Gstm1 is part of the glutathione S-transferase system, which uses reduced glutathione (GSH) for H_2_O_2_ reduction and detoxification of xenobiotic substances. Gpx1 and Gpx4 similarly use GSH to reduce H_2_O_2_ and other oxidant species, such as peroxynitrite and lipid peroxides. HO-1 may also confer antioxidant defense through multiple mechanisms.^49^ HO-1 catalyzes the reaction of heme to carbon monoxide (CO) and biliverdin, subsequently biliverdin is converted to bilirubin by biliverdin reductase. Biliverdin and bilirubin both possess powerful oxidant scavenging capacity. HO-1 derived bilirubin has also been shown to have Nox inhibitory potential.^50^ CO, similar to H_2_S, is a gaseous signaling molecule and vasodilator which has been associated with erectogenic capabilities.^51^ CO has also been shown to have anti-apoptotic effects and to provide cytoprotection against pro-inflammatory cytokines.^49^ Finally, CO has been shown to stimulate Gclc transcription.^52^ Transcription of Hmox1, the investigated genes related to glutathione synthesis and activity, as well as several other genes related to antioxidant capacity are in part regulated by the transcription factor nuclear factor E2-related factor 2 (Nrf2).^53^ H_2_S is indeed well known for its potential to induce Nrf2 activity.^54^ While there were no major influences of the WD on these cytoprotective genes/proteins, the increased ROS burden is likely to create a strain on the glutathione system. Augmentation of several of the elements of the glutathione system via H_2_S therapy will likely restore the glutathione system to strengthen the combat against this burden.

Nqo1 is another cytoprotective gene that is in part regulated by Nrf2.^55^ Nqo1 has multiple functions related to redox control, including detoxification of quinones, NAD^+^ regeneration, stabilization of proteins against proteasomal degradation, and superoxide reductase activity.^55^ The decrease in Nqo1 in the CSE^-/-^ cavernosum and the stimulation of Nqo1 by SG1002 may help to explain the protective effects of H_2_S against penile superoxide observed in this study.

We observed a significant decrease in Sod1 gene expression in CSE^-/-^ cavernosum that did not hold at the protein level, while we did not observe any other effects of CSE^-/-^ or H_2_S therapy on expressions of any of the superoxide dismutases. Similarly, we observed a stimulatory effect of SG1002 on Sirt3 gene expression that did not translate to the protein level, with no other effects of CSE^-/-^ or SG1002 on Sirt1 or Sirt3 expressions in the cavernosum. The sirtuins are a family of enzymes that regulate metabolism and transcriptional activity through histone deacetylation among other post-translational modifications, with Sirt1 and Sirt3 being most attributed to protection against oxidative stress.^15^ Some of the antioxidative effects of H_2_S in various cell systems and disease models have been attributed to induction of Sirt1 and Sirt3.^15,56^ Based on our results, we cannot conclude that H_2_S has a major influence on Sirt1 or Sirt3 in the corpus cavernosum of these models.

Similar to the glutathione antioxidant system, the thioredoxin system exerts control over the cellular redox state by reducing disulfide bonds of oxidized proteins.^57^ Thioredoxin 1 (Txn1) is a cytosolic and extracellular enzyme, whereas thioredoxin 2 (Txn2) resides in the mitochondria. Txn1 and Txn2 are maintained in their reduced state by their respective reductases (TxnRd1 and TxnRd2), which also catalyze the reductive activity of the thioredoxin systems. These thioredoxin systems may be inhibited endogenously by the thioredoxin interacting protein (Txnip).^57^ Indeed, Txnip overexpression has been found to induce oxidative stress and endothelial dysfunction in the aorta.^58^ H_2_S has previously been linked to thioredoxin activation, largely through a suppressive effect on Txnip. Similar to our finding of elevated Txnip expression in the CSE^-/-^ cavernosum, Txnip expression has been found to be elevated in the CSE^-/-^ aorta.^58^ As we observed elevated Txnip expression in the cavernosum following the Western diet that was suppressed by SG1002 treatment, Yue et al. found elevated Txnip expression in the atherosclerotic aorta which was partially blunted by treatment with a fast-releasing H_2_S donor.^59^ Interestingly, Nicholson et al. found that cardiac specific genetic depletion of Txn1 negated the protective effects H_2_S therapy on cardiac performance following myocardial infarction.^60^ The peroxiredoxins directly reduce H_2_O_2_ and peroxynitrite, although this activity relies on a reduced thioredoxin system.^61^ One particular concern of elevated superoxide levels is the quenching of NO bioavailability in the formation of peroxynitrite, which may exacerbate ED complications. Peroxynitrite may further exacerbate disease pathology through lipid peroxidation, protein oxidation, protein nitration, and loss of mitochondrial membrane potential.^62^ We investigated the mitochondrial peroxiredoxins, Prdx3 and Prdx5, both of which were decreased in the CSE^-/-^ cavernosum, which may contribute to ED through increasing mitochondrial oxidative stress, mitochondrial ROS emission, and loss of mitochondrial membrane potential. However, only Prdx3 was stimulated by SG1002 treatment, revealing a potentially complicated interplay between H_2_S and Prdx5. We are unaware of any prior research directly linking H_2_S and these peroxiredoxins, although both Prdx3 and Prdx5 transcription have been found to be controlled by the Nrf2 transcription factor.^63^

While CSE protein expression following the WD was approximately 66% that of the control levels, the molecular signature of the cavernosum following the long-term WD did not mimic that of the CSE^-/-^ condition. While impaired erectile function and elevated total interstitial ROS levels were common features, the respective molecular profiles investigated point more toward an elevation of oxidant production without a compensatory increase in antioxidant proteins following the WD, however complete loss of CSE did not result in an increase in oxidant sources, but rather a diminished antioxidant protein profile. Elevated Txnip expression was the only major molecular commonality between these two conditions. Nevertheless, our results indicate a stimulation of several antioxidant proteins in the cavernosum by the H_2_S donor that may ultimately help to reduce the total ROS burden despite the pro-oxidant conditions. This study had several limitations. Performing histology on penile tissues of these mice was outside the scope of the project, thus we were unable to determine if H_2_S influenced cavernous morphology or fibrosis. Our microdialysis method for ROS detection samples from the interstitium. Thus, we are unable to clearly define the relative contributions of each cellular compartment to the signal that we obtain. For example, it is not clear how much mitochondrial ROS may influence this signal, while the mitochondrial genes TxnRd2, Prdx3, and Prdx5 were influenced by H_2_S to some degree. Recent research indicates that mitochondrial ROS emission stimulates relaxation of penile arteries, an effect that is augmented with obesity.^64^ In this case, mitochondrial ROS may help to preserve some degree of erectile function. Moreover, our data reflects expressions at the gene and protein levels, which do not always translate to enzyme activity levels. Several post-translational modifications may influence enzyme activity. Indeed, emerging research indicates that much of the physiological influence of H_2_S may ultimately be propagated through cysteine sulfhydration.^35^ Indeed, some of the antioxidant effects of H_2_S have also been attributed to cysteine sulfhydration of various antioxidant proteins.^65–67^ Finally, although we designed the dosing of SG1002 to be identical to prior studies that have demonstrated a clear increase in circulating free sulfide and sulfane sulfur levels with SG1002 administration,^36,37^ we were unable to perform such measures for this study.

## 5. CONSLUSIONS

In conclusion, H_2_S appears to play an important role in the maintenance of erectile health and penile free radical balance. H_2_S appears to stimulate gene and protein expression of several antioxidant mediators in the corpus cavernosum, particularly those related to the function and restoration of the glutathione and thioredoxin antioxidant systems.

## CRediT authorship contribution statement

T.A.A.: Investigation, Formal Analysis, Writing – original draft preparation. P.R.R.: Investigation, Formal Analysis, Data curation. C.J.P.: Investigation, Formal Analysis, Project administration, Supervision. C.M.I.: Investigation, Formal Analysis. J.D.L.: Conceptualization, Funding acquisition, Investigation, Visualization, Supervision, Writing – original draft, Writing – review & editing.

## Data availability statement

Individual data points supporting the findings of this study are presented in the results section. Further details can be obtained upon reasonable request from the corresponding author.

## Acknowledgments

The authors thank Drs. Solomon Snyder and Bindu Paul for sharing the cystathionine γ-lyase knockout mouse model. We thank Dr. Daniel Berkowitz and Sandeep Jandu for expert technical assistance.

## REFERENCES

1. Popkin BM. Relationship between shifts in food system dynamics and acceleration of the global nutrition transition. Nutr Rev. 2017;75(2):73–82. doi:10.1093/NUTRIT/NUW064

2. Ding EL, Malik VS. Convergence of obesity and high glycemic diet on compounding diabetes and cardiovascular risks in modernizing China: An emerging public health dilemma. Global Health. 2008;4. doi:10.1186/1744-8603-4-4

3. Mariamenatu AH, Abdu EM. Overconsumption of Omega-6 Polyunsaturated Fatty Acids (PUFAs) versus Deficiency of Omega-3 PUFAs in Modern-Day Diets: The Disturbing Factor for Their “Balanced Antagonistic Metabolic Functions” in the Human Body. J Lipids. 2021;2021(1):8848161. doi:10.1155/2021/8848161

4. Wei Y, Meng Y, Li N, Wang Q, Chen L. The effects of low-ratio n-6/n-3 PUFA on biomarkers of inflammation: a systematic review and meta-analysis. Food Funct. 2021;12(1):30–40. doi:10.1039/D0FO01976C

5. Esposito K, Giugliano F, Maiorino MI, Giugliano D. Dietary Factors, Mediterranean Diet and Erectile Dysfunction. Journal of Sexual Medicine. 2010;7(7). doi:10.1111/j.1743-6109.2010.01842.x

6. Esposito K, Ciotola M, Giugliano F, et al. Mediterranean diet improves erectile function in subjects with the metabolic syndrome. Int J Impot Res. 2006;18(4). doi:10.1038/sj.ijir.3901447

7. Cao S, Hu X, Shao Y, et al. Relationship between weight-adjusted-waist index and erectile dysfunction in the United State: results from NHANES 2001-2004. Front Endocrinol (Lausanne). 2023;14:1128076. doi:10.3389/FENDO.2023.1128076/BIBTEX

8. Fillo J, Levcikova M, Ondrusova M, Breza J, Labas P. Importance of Different Grades of Abdominal Obesity on Testosterone Level, Erectile Dysfunction, and Clinical Coincidence. Am J Mens Health. 2017;11(2):240–245. doi:10.1177/1557988316642213/ASSET/IMAGES/LARGE/10.1177_1557988316642213-FIG4.JPEG

9. La Favor JD, Pierre CJ, Bivalacqua TJ, Burnett AL. Rapamycin Suppresses Penile NADPH Oxidase Activity to Preserve Erectile Function in Mice Fed a Western Diet. Biomedicines. 2021;10(1). doi:10.3390/BIOMEDICINES10010068

10. La Favor JD, Anderson EJ, Hickner RC, Wingard CJ. Erectile Dysfunction Precedes Coronary Artery Endothelial Dysfunction in Rats Fed a High-Fat, High-Sucrose, Western Pattern Diet. J Sex Med. 2013;10(3):694–703. doi:10.1111/jsm.12001

11. La Favor JD, Anderson EJ, Dawkins JT, Hickner RC, Wingard CJ. Exercise prevents Western diet-associated erectile dysfunction and coronary artery endothelial dysfunction: response to acute apocynin and sepiapterin treatment. American Journal of Physiology-Regulatory, Integrative and Comparative Physiology. 2013;305(4):R423–R434. doi:10.1152/ajpregu.00049.2013

12. Agarwal A, Nandipati KC, Sharma RK, Zippe CD, Raina R. Role of oxidative stress in the pathophysiological mechanism of erectile dysfunction. J Androl. 2006;27(3):335–347. doi:10.2164/jandrol.05136

13. Citi V, Martelli A, Gorica E, Brogi S, Testai L, Calderone V. Role of hydrogen sulfide in endothelial dysfunction: Pathophysiology and therapeutic approaches. J Adv Res. 2021;27:99–113. doi:10.1016/j.jare.2020.05.015

14. Geng B, Chang L, Pan C, et al. Endogenous hydrogen sulfide regulation of myocardial injury induced by isoproterenol. Biochem Biophys Res Commun. 2004;318(3):756–763. doi:10.1016/j.bbrc.2004.04.094

15. Corsello T, Komaravelli N, Casola A. Role of Hydrogen Sulfide in NRF2- and Sirtuin-Dependent Maintenance of Cellular Redox Balance. Antioxidants (Basel). 2018;7(10). doi:10.3390/ANTIOX7100129

16. Szabo C, Papapetropoulos A. International union of basic and clinical pharmacology. CII: Pharmacological modulation of H2S levels: H2s donors and H2S biosynthesis inhibitors. Pharmacol Rev. 2017;69(4):497–564. doi:10.1124/pr.117.014050

17. Wintner EA, Deckwerth TL, Langston W, et al. A monobromobimane-based assay to measure the pharmacokinetic profile of reactive sulphide species in blood. Br J Pharmacol. 2010;160(4):941–957. doi:10.1111/j.1476-5381.2010.00704.x

18. Gojon G, Morales GA. SG 1002 and catenated divalent organic sulfur compounds as promising hydrogen sulfide prodrugs. Antioxid Redox Signal. 2020;33(14):1010-1045. doi:10.1089/ars.2020.8060

19. Polhemus DJ, Li Z, Pattillo CB, et al. A Novel Hydrogen Sulfide Prodrug, SG1002, Promotes Hydrogen Sulfide and Nitric Oxide Bioavailability in Heart Failure Patients. Cardiovasc Ther. 2015;33(4):216–226. doi:10.1111/1755-5922.12128

20. Martinez AM, Sordia-Hernández LH, Morales JA, et al. A randomized clinical study assessing the effects of the antioxidants, resveratrol or SG1002, a hydrogen sulfide prodrug, on idiopathic oligoasthenozoospermia. Asian Pacific Journal of Reproduction. 2015;4(2):106–111. doi:10.1016/S2305-0500(15)30005-1

21. Yang G, Wu L, Jiang B, et al. H2S as a physiologic vasorelaxant: hypertension in mice with deletion of cystathionine gamma-lyase. Science. 2008;322(5901):587-590. doi:10.1126/SCIENCE.1162667

22. La Favor JD, Fu Z, Venkatraman V, Bivalacqua TJ, Van Eyk JE, Burnett AL. Molecular Profile of Priapism Associated with Low Nitric Oxide Bioavailability. J Proteome Res. 2018;17(3):1031–1040. doi:10.1021/acs.jproteome.7b00657

23. La Favor JD, Burnett AL. A microdialysis method to measure in vivo hydrogen peroxide and superoxide in various rodent tissues. Methods. 2016;109:131–140. doi:10.1016/j.ymeth.2016.07.012

24. Karakus S, Musicki B, La Favor JD, Burnett AL. cAMP-dependent post-translational modification of neuronal nitric oxide synthase neuroprotects penile erection in rats. BJU Int. 2017;120(6):861–872. doi:10.1111/bju.13981

25. Mohanty JG, Jaffe JS, Schulman ES, Raible DG. A highly sensitive fluorescent micro-assay of H2O2 release from activated human leukocytes using a dihydroxyphenoxazine derivative. J Immunol Methods. 1997;202(2):133–141. doi:10.1016/S0022-1759(96)00244-X

26. Krumschnabel G, Fontana-Ayoub M, Sumbalova Z, et al. Simultaneous high-resolution measurement of mitochondrial respiration and hydrogen peroxide production. Methods Mol Biol. 2015;1264:245–261. doi:10.1007/978-1-4939-2257-4_22

27. Pierre CJ, Azeez TA, Rossetti ML, Gordon BS, La Favor JD. Long-term administration of resveratrol and MitoQ stimulates cavernosum antioxidant gene expression in a mouse castration model of erectile dysfunction. Life Sci. 2022;310. doi:10.1016/J.LFS.2022.121082

28. Livak KJ, Schmittgen TD. Analysis of Relative Gene Expression Data Using Real-Time Quantitative PCR and the 2−ΔΔCT Method. Methods. 2001;25(4):402–408. doi:10.1006/METH.2001.1262

29. Kuo MM, Kim DH, Jandu S, et al. MPST but not CSE is the primary regulator of hydrogen sulfide production and function in the coronary artery. Am J Physiol Heart Circ Physiol. 2016;310(1):H71–H79. doi:10.1152/ajpheart.00574.2014

30. Esposito K, Giugliano D. Obesity, the metabolic syndrome, and sexual dysfunction. International Journal of Impotence Research 2005 17:5. 2005;17(5):391-398. doi:10.1038/sj.ijir.3901333

31. Souverein PC, Egberts ACG, Meuleman EJH, Urquhart J, Leufkens HGM. Incidence and determinants of sildenafil (dis)continuation: The Dutch cohort of sildenafil users. Int J Impot Res. 2002;14(4):259–265. doi:10.1038/sj.ijir.3900883

32. Moreland RB, Goldstein I, Kim NN, Traish A. Sildenafil citrate, a selective phosphodiesterase type 5 inhibitor: Research and clinical implications in erectile dysfunction. Trends in Endocrinology and Metabolism. 1999;10(3):97–104. doi:10.1016/S1043-2760(98)00127-1

33. Dadkhah F, Safarinejad MR, Asgari MA, Hosseini SY, Lashay A, Amini E. Atorvastatin improves the response to sildenafil in hypercholesterolemic men with erectile dysfunction not initially responsive to sildenafil. Int J Impot Res. 2010;22(1):51–60. doi:10.1038/ijir.2009.48

34. La Fuente JM, Sevilleja-Ortiz A, García-Rojo E, et al. Erectile dysfunction is associated with defective L-cysteine/hydrogen sulfide pathway in human corpus cavernosum and penile arteries. Eur J Pharmacol. 2020;884:173370. doi:10.1016/J.EJPHAR.2020.173370

35. Bibli SI, Hu J, Looso M, et al. Mapping the Endothelial Cell S-Sulfhydrome Highlights the Crucial Role of Integrin Sulfhydration in Vascular Function. Circulation. 2021;143(9):935–948. doi:10.1161/CIRCULATIONAHA.120.051877

36. Barr LA, Shimizu Y, Lambert JP, Nicholson CK, Calvert JW. Hydrogen sulfide attenuates high fat diet-induced cardiac dysfunction via the suppression of endoplasmic reticulum stress. Nitric Oxide. 2015;46:145–156. doi:10.1016/j.niox.2014.12.013

37. Kondo K, Bhushan S, King AL, et al. H2S protects against pressure overload-induced heart failure via upregulation of endothelial nitric oxide synthase. Circulation. 2013;127(10):1116–1127. doi:10.1161/CIRCULATIONAHA.112.000855

38. Angulo J, Peiró C, Cuevas P, et al. The novel antioxidant, AC3056 (2,6-di-t-butyl-4-((Dimethyl-4-Methoxyphenylsilyl)Methyloxy)Phenol), reverses erectile dysfunction in diabetic rats and improves NO-mediated responses in penile tissue from diabetic men. Journal of Sexual Medicine. 2009;6(2). doi:10.1111/j.1743-6109.2008.01088.x

39. Bivalacqua TJ, Armstrong JS, Biggerstaff J, et al. Gene transfer of extracellular SOD to the penis reduces O2-· and improves erectile function in aged rats. Am J Physiol Heart Circ Physiol. 2003;284(4 53-4). doi:10.1152/ajpheart.00770.2002

40. Xi Y, Feng Z, Xia T, et al. Caveolin-1 scaffolding domain-derived peptide enhances erectile function by regulating oxidative stress, mitochondrial dysfunction, and apoptosis of corpus cavernosum smooth muscle cells in rats with cavernous nerve injury. Life Sci. 2024;348. doi:10.1016/J.LFS.2024.122694

41. Andrade MR, Azeez TA, Montgomery MM, et al. Neurovascular dysfunction associated with erectile dysfunction persists after long-term recovery from simulations of weightlessness and deep space irradiation. FASEB J. 2023;37(12). doi:10.1096/FJ.202300506RR

42. Watts M, Kolluru GK, Dherange P, et al. Decreased bioavailability of hydrogen sulfide links vascular endothelium and atrial remodeling in atrial fibrillation. Redox Biol. 2021;38. doi:10.1016/J.REDOX.2020.101817

43. Barrow K, Wang Y, Yu R, Zhu J, Yang G. H2S protects from oxidative stress-driven ACE2 expression and cardiac aging. Mol Cell Biochem. 2022;477(5):1393–1403. doi:10.1007/S11010-022-04386-4

44. Xu M, Liu X, Bao P, et al. Skeletal Muscle CSE Deficiency Leads to Insulin Resistance in Mice. Antioxidants (Basel). 2022;11(11). doi:10.3390/ANTIOX11112216

45. Bryan S, Yang G, Wang R, Khaper N. Cystathionine gamma-lyase-deficient smooth muscle cells exhibit redox imbalance and apoptosis under hypoxic stress conditions. Exp Clin Cardiol. 2011;16(4):e36. Accessed January 22, 2025. https://pmc.ncbi.nlm.nih.gov/articles/PMC3206107/

46. Ng HH, Yildiz GS, Ku JM, Miller AA, Woodman OL, Hart JL. Chronic NaHS treatment decreases oxidative stress and improves endothelial function in diabetic mice. Diab Vasc Dis Res. 2017;14(3):246–253. doi:10.1177/1479164117692766

47. Xiao L, Dong JH, Jin S, et al. Hydrogen Sulfide Improves Endothelial Dysfunction via Downregulating BMP4/COX-2 Pathway in Rats with Hypertension. Oxid Med Cell Longev. 2016;2016. doi:10.1155/2016/8128957

48. Zhang J, Cai X, Zhang Q, et al. Hydrogen sulfide restores sevoflurane postconditioning mediated cardioprotection in diabetic rats: Role of SIRT1/Nrf2 signaling-modulated mitochondrial dysfunction and oxidative stress. J Cell Physiol. 2021;236(7):5052–5068. doi:10.1002/JCP.30214

49. Consoli V, Sorrenti V, Grosso S, Vanella L. Heme Oxygenase-1 Signaling and Redox Homeostasis in Physiopathological Conditions. Biomolecules. 2021;11(4). doi:10.3390/BIOM11040589

50. Mazza F, Goodman A, Lombardo G, Vanella A, Abraham NG. Heme oxygenase-1 gene expression attenuates angiotensin II-mediated DNA damage in endothelial cells. Exp Biol Med (Maywood). 2003;228(5):576–583. doi:10.1177/15353702-0322805-31

51. Abdel Aziz MT, Mostafa T, Atta H, et al. Putative role of carbon monoxide signaling pathway in penile erectile function. J Sex Med. 2009;6(1):49–60. doi:10.1111/J.1743-6109.2008.01050.X

52. Li MH, Jang JH, Na HK, Cha YN, Surh YJ. Carbon Monoxide Produced by Heme Oxygenase-1 in Response to Nitrosative Stress Induces Expression of Glutamate-Cysteine Ligase in PC12 Cells via Activation of Phosphatidylinositol 3-Kinase and Nrf2 Signaling. Journal of Biological Chemistry. 2007;282(39):28577–28586. doi:10.1074/JBC.M701916200

53. Tonelli C, Chio IIC, Tuveson DA. Transcriptional Regulation by Nrf2. Antioxid Redox Signal. 2018;29(17):1727–1745. doi:10.1089/ars.2017.7342

54. Yang G, Zhao K, Ju Y, et al. Hydrogen sulfide protects against cellular senescence via s-sulfhydration of keap1 and activation of Nrf2. Antioxid Redox Signal. 2013;18(15):1906–1919. doi:10.1089/ars.2012.4645

55. Ross D, Siegel D. The diverse functionality of NQO1 and its roles in redox control. Redox Biol. 2021;41. doi:10.1016/J.REDOX.2021.101950

56. Xie L, Feng H, Li S, et al. SIRT3 Mediates the Antioxidant Effect of Hydrogen Sulfide in Endothelial Cells. Antioxid Redox Signal. 2016;24(6):329–343. doi:10.1089/ARS.2015.6331

57. Lu J, Holmgren A. The thioredoxin antioxidant system. Free Radic Biol Med. 2014;66:75–87. doi:10.1016/J.FREERADBIOMED.2013.07.036

58. Tian D, Dong J, Jin S, Teng X, Wu Y. Endogenous hydrogen sulfide-mediated MAPK inhibition preserves endothelial function through TXNIP signaling. Free Radic Biol Med. 2017;110:291–299. doi:10.1016/J.FREERADBIOMED.2017.06.016

59. Yue LM, Gao YM, Han BH. Evaluation on the effect of hydrogen sulfide on the NLRP3 signaling pathway and its involvement in the pathogenesis of atherosclerosis. J Cell Biochem. 2019;120(1):481–492. doi:10.1002/JCB.27404

60. Nicholson CK, Lambert JP, Molkentin JD, Sadoshima J, Calvert JW. Thioredoxin 1 is essential for sodium sulfide-mediated cardioprotection in the setting of heart failure. Arterioscler Thromb Vasc Biol. 2013;33(4):744–751. doi:10.1161/ATVBAHA.112.300484

61. Kameritsch P, Singer M, Nuernbergk C, et al. The mitochondrial thioredoxin reductase system (TrxR2) in vascular endothelium controls peroxynitrite levels and tissue integrity. Proc Natl Acad Sci U S A. 2021;118(7). doi:10.1073/PNAS.1921828118

62. Pacher P, Beckman JS, Liaudet L. Nitric oxide and peroxynitrite in health and disease. Physiol Rev. 2007;87(1):315–424. doi:10.1152/PHYSREV.00029.2006/ASSET/IMAGES/LARGE/Z9J0010724240018.JPEG

63. Miyamoto N, Izumi H, Miyamoto R, et al. Quercetin induces the expression of peroxiredoxins 3 and 5 via the Nrf2/NRF1 transcription pathway. Invest Ophthalmol Vis Sci. 2011;52(2):1055–1063. doi:10.1167/IOVS.10-5777

64. Gómez del Val A, Sánchez A, Freire-Agulleiro Ó, et al. Penile endothelial dysfunction, impaired redox metabolism and blunted mitochondrial bioenergetics in diet-induced obesity: Compensatory role of H2O2. Free Radic Biol Med. 2025;230:222-233. doi:10.1016/J.FREERADBIOMED.2025.02.004

65. Cheung SH, Lau JYW. Hydrogen sulfide mediates athero-protection against oxidative stress via S-sulfhydration. PLoS One. 2018;13(3):e0194176. doi:10.1371/JOURNAL.PONE.0194176

66. Du C, Lin X, Xu W, et al. Sulfhydrated Sirtuin-1 Increasing Its Deacetylation Activity Is an Essential Epigenetics Mechanism of Anti-Atherogenesis by Hydrogen Sulfide. https://home.liebertpub.com/ars. 2018;30(2):184-197. doi:10.1089/ARS.2017.7195

67. Xie ZZ, Shi MM, Xie L, et al. Sulfhydration of p66Shc at Cysteine59 mediates the antioxidant effect of hydrogen sulfide. Antioxid Redox Signal. 2014;21(18):2531–2542. doi:10.1089/ARS.2013.5604/ASSET/IMAGES/LARGE/FIGURE7.JPEG

