## Supplemental Data for "Hydrogen sulfide protects erectile function through stimulation of antioxidant defense in the corpus cavernosum"

| Gene | Description | Assay ID | Amplicon Size | Accession Number |
|---|---|---|---|---|
| Cth | Cystathionine gamma-lyase | Mm00461247_m1 | 67 | NM_145953.2 |
| Cbs | Cystathionine beta-synthase | Mm00460654_m1 | 125 | NM_178224.3 |
| Mpst | Mercaptopyruvate sulfurtranserase | Mm00460389_m1 | 67 | NM_138670.3 |
| Nqo1 | NADPH quinone dehydrogenase 1 | Mm01253561_m1 | 81 | NM_008706.5 |
| Gclc | Glutamate-Cysteine Ligase Catalytic Subunit | Mm00802658_m1 | 78 | NM_010295.2 |
| Gclm | Glutamate-Cysteine Ligase Modifier Subunit | Mm00514996_m1 | 76 | NM_008129.4 |
| Gstm1 | Glutathione S-Transferase 1 | Mm00833915_g1 | 177 | NM_010358.5 |
| Gpx1 | Glutathione Peroxidase 1 | Mm00656767_g1 | 134 | NM_008160.6 |
| Gpx4 | Glutathione Peroxidase 4 | Mm00515041_m1 | 103 | NM_008162.3 |
| Hmox1 | Heme Oxygenase 1 | Mm00516005_m1 | 69 | NM_010442.2 |
| Cat | Catalase | Mm00437992_m1 | 64 | NM_009804.2 |
| Sod1 | Superoxide Dismutase 1 | Mm01344233_g1 | 71 | NM_011434.1 |
| Sod2 | Superoxide Dismutase 2 | Mm01313000_m1 | 67 | NM_013671.3 |
| Sod3 | Superoxide Dismutase 3 | Mm00448831_s1 | 97 | NM_011435.3 |
| Sirt1 | Sirtuin 1 | Mm01168521_m1 | 94 | NM_019812.2 |
| Sirt3 | Sirtuin 3 | Mm00452131_m1 | 68 | NM_022433.2 |
| Txn1 | Thioredoxin 1 | Mm00726847_s1 | 113 | NM_011660.3 |
| Txnrd1 | Thioredoxin Reductase 1 | Mm00443675_m1 | 63 | NM_015762.2 |
| Txn2 | Thioredoxin 2 | Mm00444931_m1 | 69 | NM_019913.5 |
| TxnRd2 | Thioredoxin Reductase 2 | Mm00496766_m1 | 56 | NM_013711.3 |
| Txnip | Thioredoxin Interacting Protein | Mm01265659_g1 | 71 | NM_023719.2 |
| Prdx3 | Peroxiredoxin 3 | Mm00545848_m1 | 133 | NM_007452.2 |
| Prdx5 | Peroxiredoxin 5 | Mm00465365_m1 | 70 | NM_012021.2 |
| Hprt1 | Hypoxanthine Guanine Phosporibosyl Transferase | Mm03024075_m1 | 131 | NM_013556.2 |

Table S1. Taqman primers used for qRT-PCR.

| Protein | Description | Antibody Manufacturer | Catalog Number | Antibody Dilution |
|---|---|---|---|---|
| CSE | Cystathionine gamma-lyase | Proteintech | 12217-1-AP | 1:1500 |
| CBS | Cystathionine beta-synthase | Cell Signaling | 14782 | 1:1000 |
| MPST | Mercaptopyruvate sulfurtranserase | Sigma Aldrich | HPA001240 | 1:1000 |
| GCLC | Glutamate-Cysteine Ligase Catalytic Subunit | Proteintech | 12601-1-AP | 1:2000 |
| GPX4 | Glutathione Peroxidase 4 | Cell Signaling | 52455 | 1:1000 |
| HO-1 | Heme Oxygenase 1 | Cell Signaling | 82206 | 1:1000 |
| SOD1 | Superoxide Dismutase 1 | Proteintech | 10269-1-AP | 1:1000 |
| SOD2 | Superoxide Dismutase 2 | Cell Signaling | 13141 | 1:3000 |
| SOD3 | Superoxide Dismutase 3 | Santa Cruz | sc-271170 | 1:1000 |
| SIRT1 | Sirtuin 1 | Cell Signaling | 9475 | 1:1000 |
| SIRT3 | Sirtuin 3 | Cell Signaling | 5490 | 1:1000 |
| TXN1 | Thioredoxin 1 | Cell Signaling | 2298 | 1:1000 |
| TXNRD1 | Thioredoxin Reductase 1 | Cell Signaling | 6925 | 1:1000 |
| TXNIP | Thioredoxin Interacting Protein | Proteintech | 18243-1-AP | 1:1000 |
| PRDX3 | Peroxiredoxin 3 | Proteintech | 10664-1-AP | 1:2000 |
| PRDX5 | Peroxiredoxin 5 | Proteintech | 17724-1-AP | 1:1000 |
| Nox2 | NADPH oxidase 2 | BD | 611414 | 1:1000 |
| $p22^{phox}$ | 47 kDa NADPH oxidase subunit | Santa Cruz | sc-130550 | 1:500 |
| $p47^{phox}$ | 67 kDa NADPH oxidase subunit | Santa Cruz | sc-17845 | 1:1000 |
| $p67^{phox}$ | 22 kDa NADPH oxidase subunit | Proteintech | 15551-1-AP | 1:1000 |
| Nox4 | NADPH oxidase 4 | Invitrogen | PA5-72816 | 1:2000 |
| XO | Xanthine Oxydase | Proteintech | 55156-1-AP | 1:1000 |
| GAPDH | Glyceraldehyde-3-phosphate dehydrogenase | Proteintech | 60004-1-Ig or 10494-1-AP | 1:4000 |
| | Anti-Mouse Secondary Antibody | Cell Signaling | 7076 | 1:4000 |
| | Anti-Rabbit Secondary Antibody | Bethyl Laboratories | A120-101P | 1:10000 |

Table S2. Antibodies used for immunoblotting.

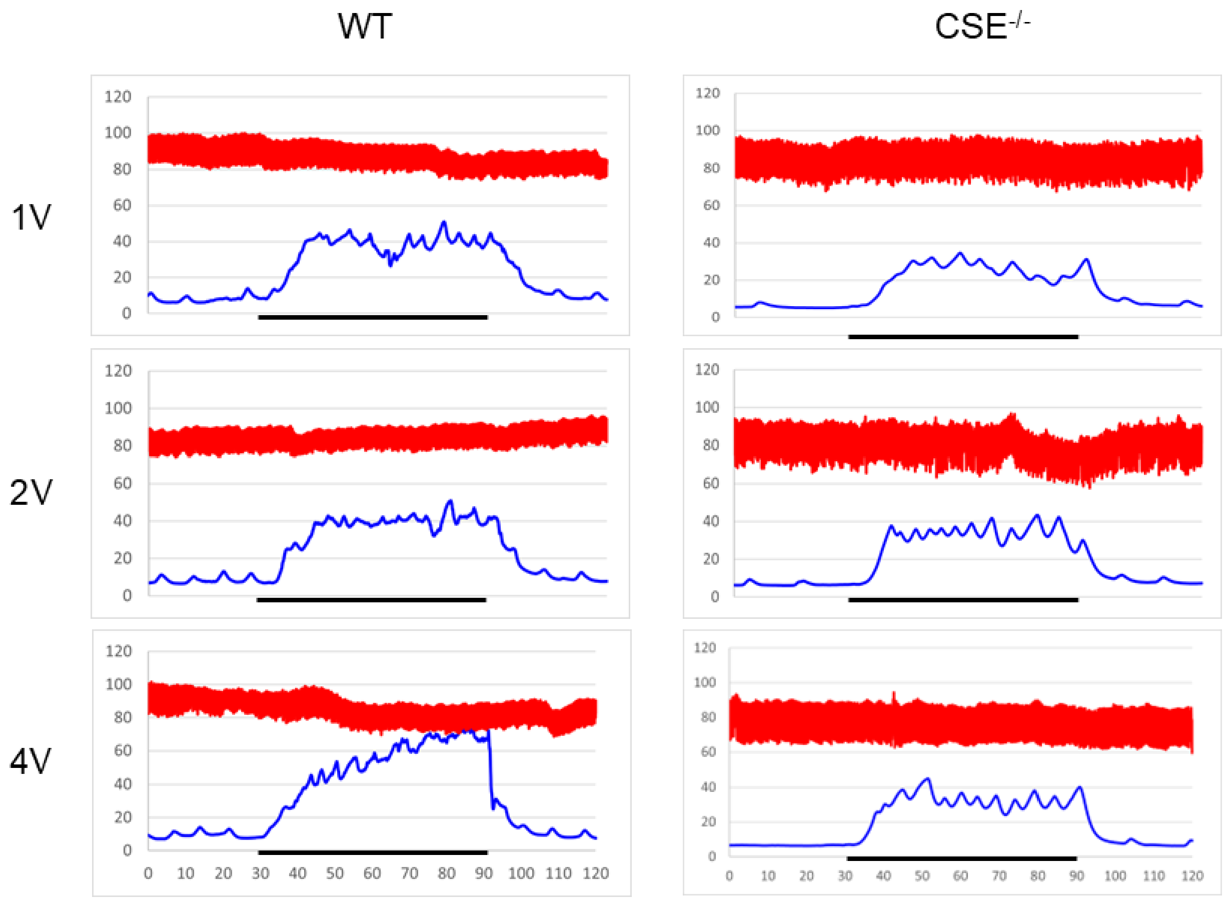


Figure S1. Representative tracings for erectile function assessment of WT and CSE$^{-/-}$ mice. Tracings are presented for 1, 2, and 4 V stimulations of the cavernous nerve. The red tracing represents mean arterial pressure (MAP) obtained from the carotid artery. The blue tracing represents intracavernous pressure (ICP). The x-axis represents time (s). The y-axis represents pressure (mmHg). The black line from 30-90 s represents the period of electrical stimulation.

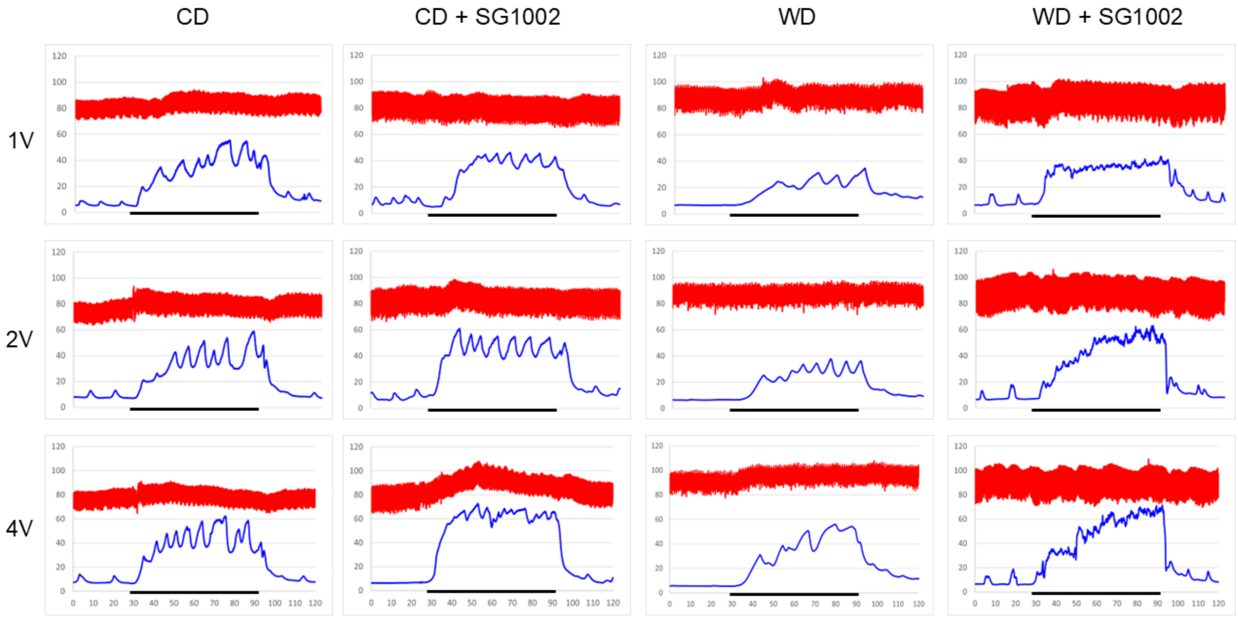


Figure S2. Representative tracings for erectile function assessment of mice following the 18-week dietary intervention. Tracings are presented for 1, 2, and 4 V stimulations of the cavernous nerve. The red tracing represents mean arterial pressure (MAP) obtained from the carotid artery. The blue tracing represents intracavernous pressure (ICP). The x-axis represents time (s). The y-axis represents pressure (mmHg). The black line from 30-90 s represents the period of electrical stimulation.

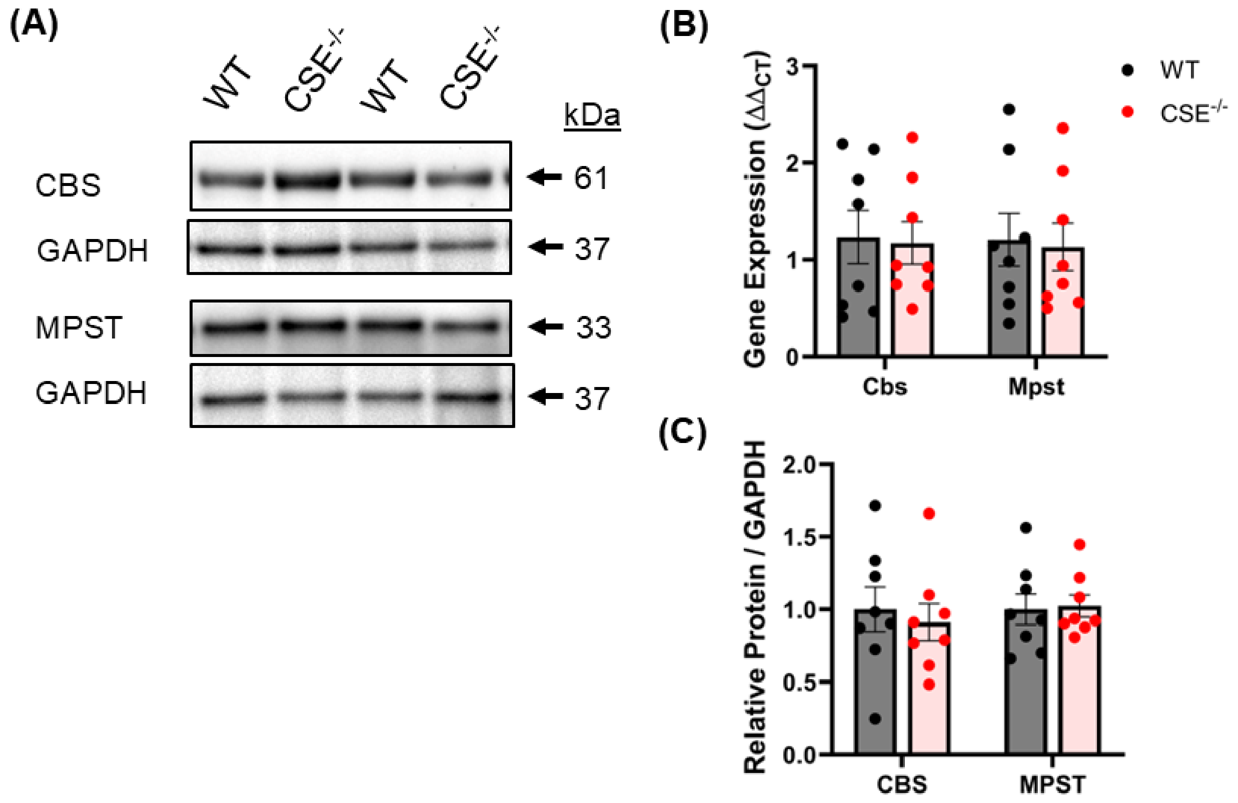


Figure S3. Cystathionine γ-lyase (CSE) deficiency does not impact corpus cavernosum gene or protein expression of the $H_2S$ producing enzymes CBS or MPST. (A) Representative images of western blots run for CBS and MPST and their respective loading control protein GAPDH. (B) Gene expression of Cbs and Mpst assessed by qRT-PCR. (C) Quantification of normalized band intensity density of proteins assessed by western blot. Bars represent mean ± SEM for n = 8/group.

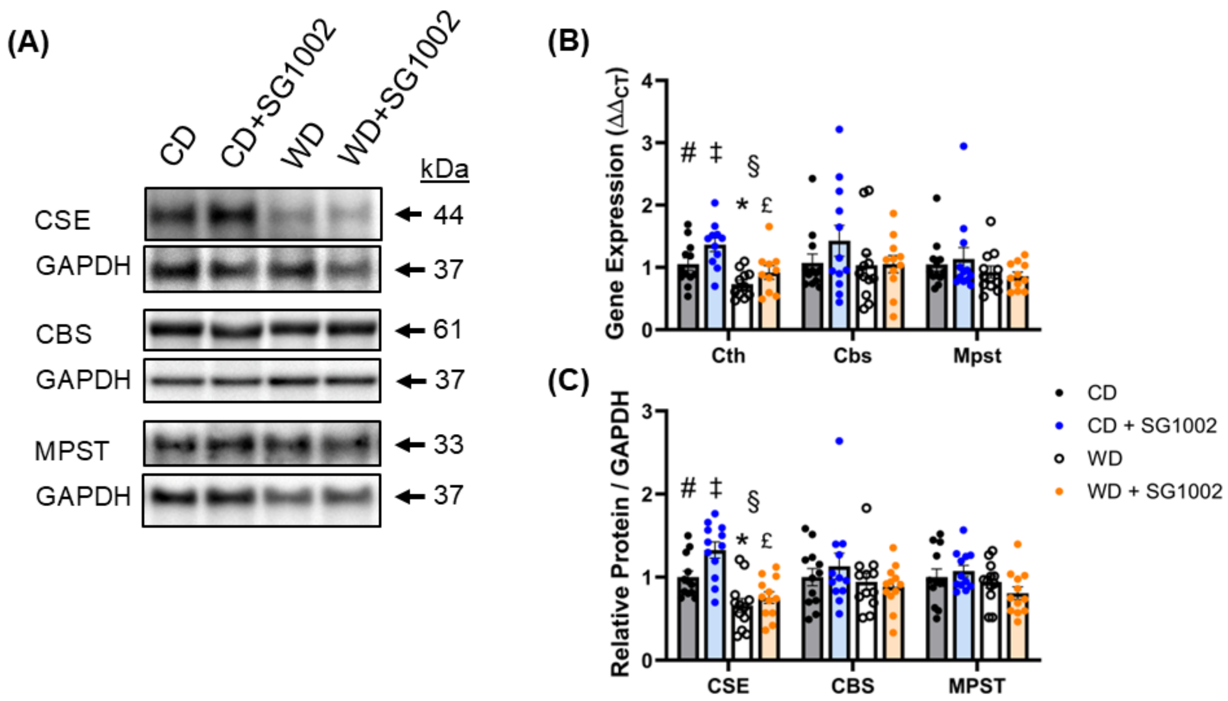


Figure S4. Effect of H2S therapy on gene and protein expression of H2S production enzymes in the corpus cavernosum. Measures taken following the 18-week control diet (CD) or Western diet (WD) with and without the $H_2S$ prodrug SG1002 for the final 6 weeks. (A) Representative images of western blots run for CSE, CBS, and MPST, as well as the respective loading control protein GAPDH. (B) Gene expression of Cth, Cbs, and Mpst assessed by qRT-PCR. (C) Quantification of normalized band intensity density of proteins assessed by western blot. Bars represent mean ± SEM for n = 10-12/group. # p<.05 two-way ANOVA main effect of SG1002 treatment, § p<.05 two-way ANOVA main effect of diet. The following indicate a significant difference between groups via post-hoc analysis: * CD vs. WD; ‡ CD vs. CD + SG1002; £ CD + SG1002 vs. WD + SG1002.

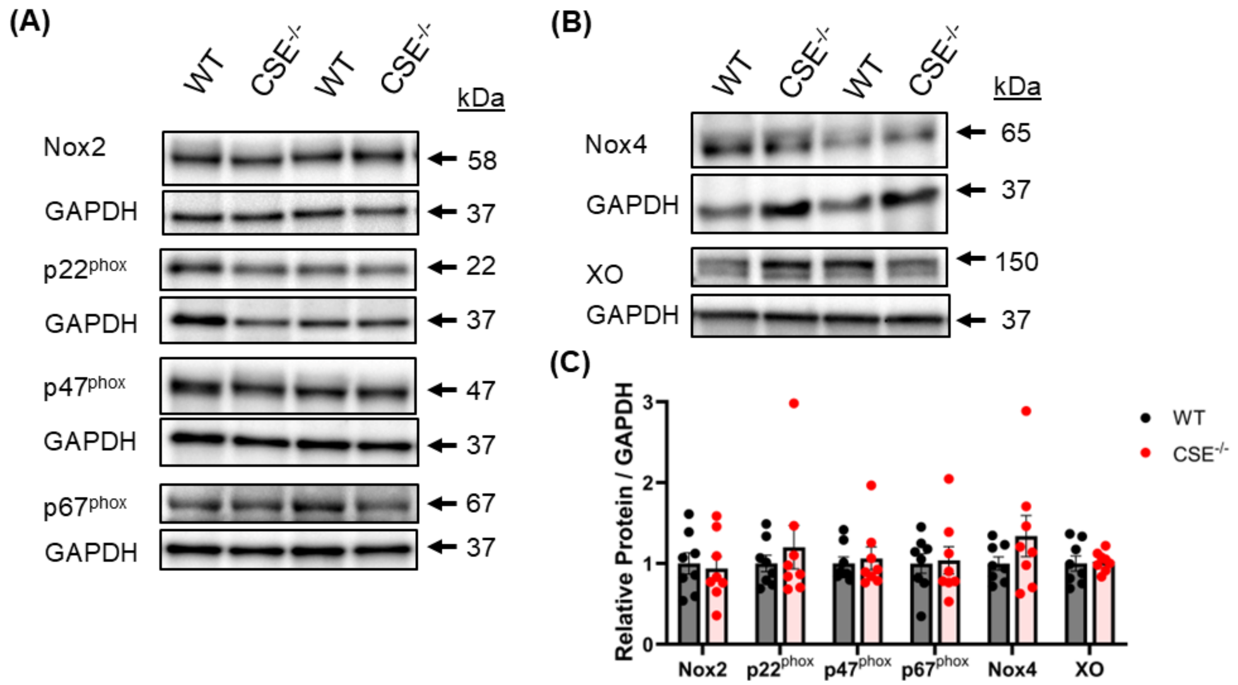


Figure S5. Cystathionine γ-lyase (CSE) deficiency does not affect protein expression of components of the NADPH oxidase or xanthine oxidase oxidant generation systems in the corpus cavernosum. (A) Representative images of western blots run for Nox2, $p22^{phox}$, $p47^{phox}$, $p67^{phox}$, (B) Nox4, and XO, as well as the respective loading control protein GAPDH. (C) Quantification of normalized band intensity density of proteins assessed by western blot. Bars represent mean ± SEM for n = 8/group.

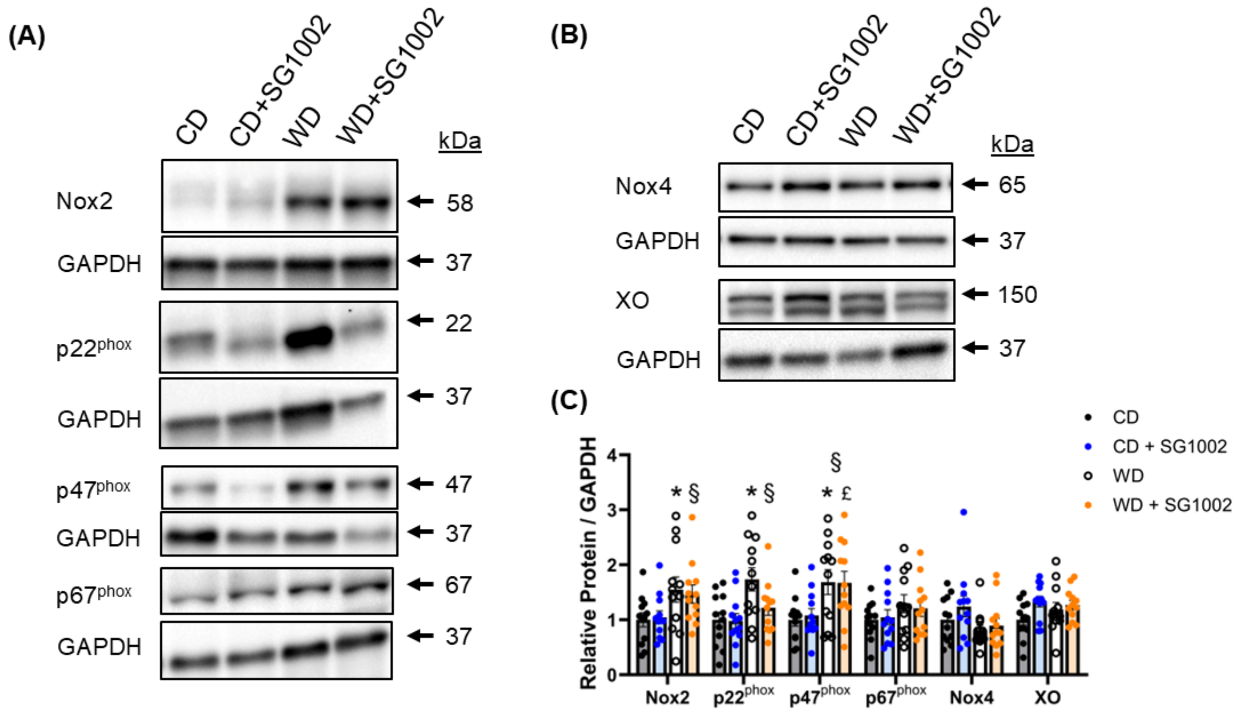


Figure S6. $H_2S$ therapy marginally impacts components of the NADPH oxidase or xanthine oxidase oxidant generation systems in the corpus cavernosum. (A) Representative images of western blots run for Nox2, $p22^{phox}$, $p47^{phox}$, $p67^{phox}$, (B) Nox4, and XO, as well as the respective loading control protein GAPDH. (C) Quantification of normalized band intensity density of proteins assessed by western blot. Bars represent mean ± SEM for n = 12/group. § p<.05 two-way ANOVA main effect of diet. The following indicate a significant difference between groups via post-hoc analysis: * CD vs. WD; † WD vs. WD + SG1002. £ CD + SG1002 vs. WD + SG1002.
